# Chemical screening of food-related chemicals for human fatty liver risk: Combining high content imaging of cellular responses with in vitro to in vivo extrapolation

**DOI:** 10.1101/2022.06.02.494529

**Authors:** Fabrice A. Müller, Marianna Stamou, Felix Englert, Ole Frenzel, Sabine Diedrich, John F. Wambaugh, Shana J. Sturla

**Affiliations:** Department of Health Sciences and Technology, ETH Zurich, 8092 Zurich, Switzerland; Center for Computational Toxicology and Exposure, Office of Research and Development, United States Environmental Protection Agency, Research Triangle Park, North Carolina 27711, United States

**Author notes:** Corresponding author: Prof. Dr. Shana J. Sturla, ETH Zurich, Department of Health Sciences and Technology, Schmelzbergstrasse 9, 8092 Zurich, Switzerland.

**Keywords:** new approach method, high-content imaging, liver toxicity, in vitro to *in vivo* extrapolation

## Abstract

Nonalcoholic fatty liver disease (NAFLD) is an increasingly prevalent human disease with accumulating evidence linking its pathophysiology and co-morbidities to chemical exposures. The complex pathophysiology of NAFLD has limited the elucidation of potential chemical etiologies. In this study we generated a high-content imaging analysis method for the simultaneous quantification of sentinel steatosis cellular markers in chemically exposed human liver cells in vitro combined with a computational model for the extrapolation of human oral equivalent doses (OED). First, the in vitro test method was generated using 14 reference chemicals with known capacities to induce cellular alterations in nuclear morphology, lipid accumulation, mitochondrial membrane potential and oxidative stress. These effects were quantified on a single cell- and population-level, and then, using physiologically based pharmacokinetic modelling and reverse dosimetry, OEDs were extrapolated from these in vitro data. The extrapolated OEDs were confirmed to be within biologically relevant exposure ranges for the reference chemicals. Next, we tested 14 chemicals found in food, selected from thousands of putative chemicals on the basis of structure-based prediction for nuclear receptor activation. Amongst these, orotic acid had an extrapolated OED overlapping with realistic exposure ranges. By the strategy developed in this study, we were able to characterize known NAFLD-inducing chemicals and translate to data scarce food-related chemicals, amongst which we identified orotic acid to induce steatosis. This strategy addresses needs of next generation risk assessment, and can be used as a first chemical prioritization hazard screening step in a tiered approach to identify chemical risk factors for NAFLD.

## 2 Introduction

Nonalcoholic fatty liver disease (NAFLD) affects around 25% of the adult population and 70-80% of obese or diabetic individuals (Younossi *et al*., 2018; Chalasani *et al*., 2012). It is a complex spectrum of diseases ranging from benign liver steatosis to nonalcoholic steatohepatitis (NASH). Steatosis is a hepatic accumulation of fatty acids where the total liver fat content exceeds 5%. (Musso *et al*., 2016; Chalasani *et al*., 2012). NAFLD risk factors include metabolic disorder (obesity, type 2 diabetes), genetics, drugs, and environmental exposure to chemicals (Wahlang *et al*., 2013; Kaiser *et al*., 2012). A number of drugs, such as amiodarone, valproic acid, and cancer chemotherapeutics cause NAFLD in some patients, particularly after chronic treatment (Begriche *et al*., 2011; Schumacher and Guo, 2015; Willebrords *et al*., 2015; Jennings *et al*., 2014). Exposure to environmental chemicals, such as solvents, persistent organic pollutants, and pesticides, has been suggested also to be a risk factor (Wahlang *et al*., 2013; Kaiser *et al*., 2012). In addition, its link to obesity and metabolic disorders suggest food-related exposures could contribute to NAFLD risk, but these have not been systematically assessed for their capacity to induce steatosis or other sentinel molecular processes associated with NAFLD risk.

The adverse outcome pathway (AOP) for liver steatosis involves binding to a nuclear receptor as the molecular initiating event, which leads to triglyceride accumulation, mitochondrial disruption and nucleus distortion (Vinken, 2015). Nuclear receptor activation has been associated with drug-induced liver injury, and recently a number of nuclear receptors have been linked to the onset of hepatic steatosis. Their activation can lead to the inhibition of β-oxidation and an increase of de novo fatty acid synthesis which creates an imbalance in the fatty acid metabolism and subsequently liver triglycerides accumulate. Ongoing accumulation can induce nucleus distortion, further disruption of mitochondrial functionalities and endoplasmic reticulum stress eventually leading to fatty liver cells and steatosis. (Mellor *et al*., 2015; Moya *et al*., 2010). It is challenging to predict which chemicals could induce the key events and steatotic phenotype associated with NAFLD, and to determine their relative potency while minimizing *in vivo* studies.

There are at least four examples of quantitative in vitro assays for steatosis hazard assessment available (**Table 1**) (Luckert *et al*., 2018; Tolosa *et al*., 2016; Shah *et al*., 2021; Donato *et al*., 2012). In a previous study (Luckert *et al*., 2018), we characterized the molecular signature of cyproconazole-induced steatosis in a metabolically competent model of human liver cells (HepaRG) by investigating key events along the liver steatosis AOP (Mellor *et al*., 2015). This approach involved using a battery of *in vitro* assays: nuclear receptor activation, gene and protein expression, lipid accumulation and mitochondrial respiration. It involved a major collaborative effort for performing independent but highly coordinated measurements, allowing testing of only a small number of compounds at considerable cost. In another example, Tolosa and Donato developed an in vitro strategy to characterize the steatogenic potential of chemicals by using a high-content lipid accumulation imaging approach (Tolosa *et al*., 2016; Donato *et al*., 2012). This method was applied to analyze 50 pharmaceuticals consisting of 32 steatotic and 18 nonsteatotic drugs in HepaRG and HepG2 cells. A key finding was that lipid accumulation could be induced by known steatotic drugs in both cell lines and that HepaRG cells were more sensitive in identifying steatosis inducing pharmaceuticals. Finally, in a recent study Shah et al. quantified mitochondrial function, lipid accumulation, endoplasmic reticulum stress, lysosomal mass, DNA texture, nuclear size, apoptosis, and cell number using a high content imaging approach in rat hepatocytes exposed to 51 hepatotoxicants. Moreover, the *in vitro* data was extrapolated to *in vivo* administered equivalent doses and compared to *in vivo* rat data (Shah *et al*., 2021). Thus, a major benefit was simultaneous quantification of several cellular responses as well as extrapolation to in vivo doses. To our knowledge, there are as yet no reports of combining an in vitro approach for simultaneous quantification of several steatosis hallmarks in human cells with extrapolation to human exposures and application for screening chemicals with unknown potential to cause NAFLD, including food-relevant exposures.

**Table 1.** Summary of selected in vitro strategies assessing liver steatosis

| Reference | Cells | Key events | Assay format | IVIVE | Chemical(s) tested |
| --- | --- | --- | --- | --- | --- |
| (Luckert *et al.*, 2018) | HepaRG | Nuclear receptor activation<br>Gene and protein expression<br>Lipid accumulation<br>Mitochondrial respiration<br>Formation of fatty liver cells | Battery to test key events along the liver steatosis AOP | none | Cyproconazole |
| (Donato *et al.*, 2012) | HepG2 | Lipid content<br>ROS generation<br>Mitochondrial membrane potential<br>Cell viability | Simultaneous assessment of key events | none | 22 pharmaceuticals consisting of known steatotic and nonsteatotic chemicals |
| (Tolosa *et al.*, 2016) | HepaRG | Lipid content<br>Viability<br>Mitochondrial membrane potential<br>ROS production | Simultaneous assessment of key events | Comparison of *in vitro* concentration to C<sub>max</sub> values | 28 pharmaceuticals consisting of known steatotic and nonsteatotic chemicals |
| (Shah *et al.*, 2021) | Rat hepatocytes | Endoplasmic reticulum stress<br>Mitochondrial function<br>Lysosomal mass<br>Steatosis<br>Apoptosis<br>DNA texture<br>Nuclear size<br>Cell number | Simultaneous assessment of key events | Prediction of administered equivalent doses using toxicokinetic modeling and comparing to *in vivo* exposure data | 51 chemicals known to induce hepatotoxicity |

IVIVE: *in vitro* to *in vivo* extrapolation

In this study, we used high-content imaging to simultaneously quantify dose-response relationships in human liver cells for multiple cellular responses that are hallmarks of steatosis, including chemical-induced lipid accumulation, mitochondrial membrane potential disruption, oxidative stress, and nuclear morphology changes. Responses were evaluated in a co-culture of hepatocyte- and cholangiocyte-like cells (HepaRG) on both single cell and population levels. Fourteen reference chemicals with known steatotic effects by different mechanisms were tested in order to dissect distinct patterns of molecular responses, relevant alone and in combination to the etiology of NAFLD. To evaluate the exposure and physiological relevance of the dose-response relationship, we integrated the in vitro data for the reference compounds with physiologically based pharmacokinetic modeling (PBPK) and reverse dosimetry to derive human dose equivalents. These levels were then compared to published in vivo exposure values. Finally, on the basis of predicted nuclear receptor agonist potential and literature evidence for NAFLD relevance, we curated a panel of food-related compounds and pesticides, and tested these using the new approach method.

## 3 Materials and Methods

### Chemicals and reagents

All tested chemicals (**Table 2**) were purchased from Sigma-Aldrich (St. Louis, Missouri). Chemical stock solutions (200x) were prepared in dimethylsulfoxide (DMSO). Chemicals soluble in water were directly dissolved in cell culture medium. A mixture of oleic acid/palmitic acid (OA/PA) was complexed to bovine serum albumin (BSA) in a ratio of 5.5:1 (OA/PA: BSA). OA/PA was first dissolved in DMSO and then added to exposure medium supplemented with the corresponding BSA concentration. The OA/PA medium mix was then warmed at 60 °C for 1 h (to aid dissolution), allowed to return to room temperature, and subsequently used for experiments.

**Table 2.**
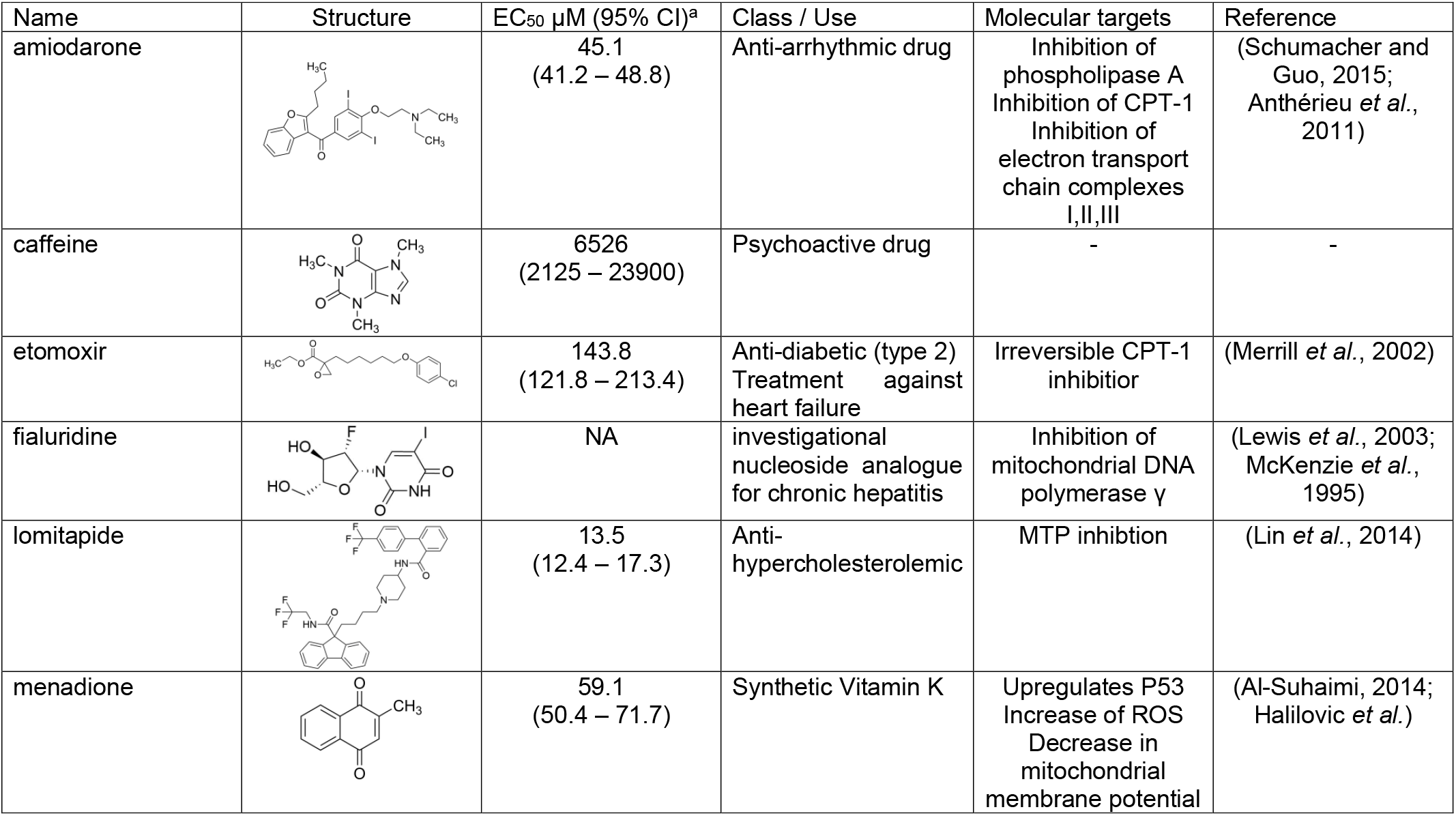
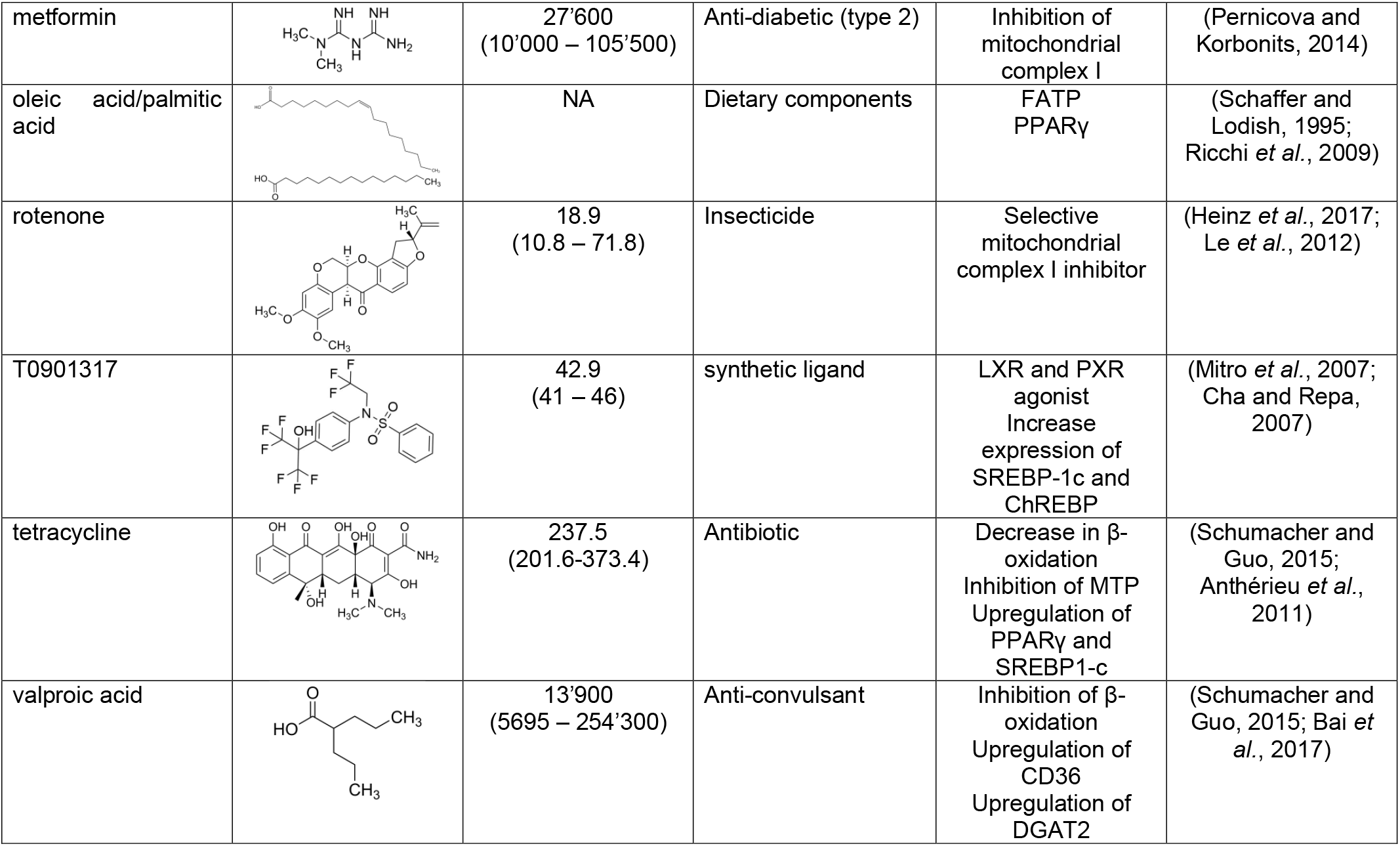
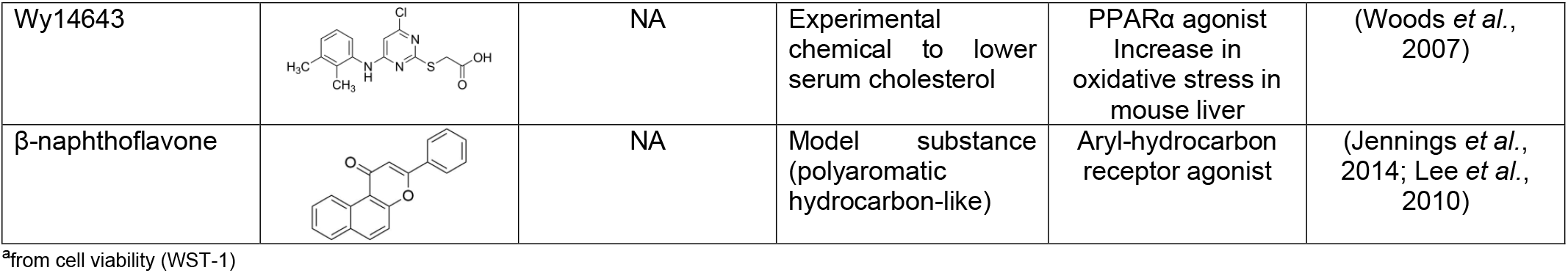
Reference chemicals used to develop the high-content screening assay.

| Name | Structure | EC<sub>50</sub> μM (95% CI)<sup>a</sup> | Class / Use | Molecular targets | Reference |
| --- | --- | --- | --- | --- | --- |
| amiodarone |  | 45.1<br>(41.2 – 48.8) | Anti-arrhythmic drug | Inhibition of phospholipase A<br>Inhibition of CPT-1<br>Inhibition of electron transport chain complexes I,II,III | (Schumacher and Guo, 2015; Anthérieu *et al.*, 2011) |
| caffeine |  | 6526<br>(2125 – 23900) | Psychoactive drug | - | - |
| etomoxir |  | 143.8<br>(121.8 – 213.4) | Anti-diabetic (type 2)<br>Treatment against heart failure | Irreversible CPT-1 inhibitor | (Merrill *et al.*, 2002) |
| fialuridine | | NA | investigational nucleoside analogue for chronic hepatitis | Inhibition of mitochondrial DNA polymerase $\gamma$ | (Lewis *et al.*, 2003; McKenzie *et al.*, 1995) |
| lomitapide |  | 13.5<br>(12.4 – 17.3) | Anti-hypercholesterolemic | MTP inhibition | (Lin *et al.*, 2014) |
| menadione |  | 59.1<br>(50.4 – 71.7) | Synthetic Vitamin K | Upregulates P53<br>Increase of ROS<br>Decrease in mitochondrial membrane potential | (Al-Suhaimi, 2014; Halilovic *et al.*) |
| metformin |  | 27'600<br>(10'000 – 105'500) | Anti-diabetic (type 2) | Inhibition of mitochondrial complex I | (Pernicova and Korbonits, 2014) |
| oleic acid/palmitic acid | | NA | Dietary components | FATP<br>PPARγ | (Schaffer and Lodish, 1995; Ricchi *et al.*, 2009) |
| rotenone |  | 18.9<br>(10.8 – 71.8) | Insecticide | Selective mitochondrial complex I inhibitor | (Heinz *et al.*, 2017; Le *et al.*, 2012) |
| T0901317 |  | 42.9<br>(41 – 46) | synthetic ligand | LXR and PXR agonist<br>Increase expression of SREBP-1c and ChREBP | (Mitro *et al.*, 2007; Cha and Repa, 2007) |
| tetracycline | | 237.5<br>(201.6-373.4) | Antibiotic | Decrease in β-oxidation<br>Inhibition of MTP<br>Upregulation of PPARγ and SREBP1-c | (Schumacher and Guo, 2015; Anthérieu *et al.*, 2011) |
| valproic acid | | 13'900<br>(5695 – 254'300) | Anti-convulsant | Inhibition of β-oxidation<br>Upregulation of CD36<br>Upregulation of DGAT2 | (Schumacher and Guo, 2015; Bai *et al.*, 2017) |
| Wy14643 | | NA | Experimental chemical to lower serum cholesterol | PPARα agonist<br>Increase in oxidative stress in mouse liver | (Woods *et al.*, 2007) |
| β-naphthoflavone | | NA | Model substance (polyaromatic hydrocarbon-like) | Aryl-hydrocarbon receptor agonist | (Jennings *et al.*, 2014; Lee *et al.*, 2010) |

<sup>a</sup>from cell viability (WST-1)

### Cell culture and chemical exposure

Undifferentiated HepaRG cells were purchased from Biopredic international (Saint Grégoire, France). Cells were seeded at a density of approximately 27’000 cells/cm^2^ into a 150 × 25 mm petri dish and were cultured for 2 weeks in William’s E medium (cat. nr.: 32551020, ThermoFisher Scientific, Massachusetts, USA) supplemented with 2 mM glutamine, 10 % (v/v) fetal bovine serum (FBS) good forte (cat. nr.: P40-47500, PAN-Biotech, Aidenbach, Germany), 100 U/ml penicillin and 100 μg/ml streptomycin (cat. nr.: 15140122, ThermoFisher Scientific, Massachusetts, USA), 5 μg/ml gibco recombinant AOF insulin (cat. nr.: A11382II, ThermoFisher Scientific, Massachusetts, USA) and 50 μM hydrocortisone hemisuccinate (cat. nr.: H4881, Sigma-Aldrich, St. Louis, USA) at 37°C and 5% CO_2_. Differentiation was induced by culturing the cells for another 2 weeks in the above mentioned medium supplemented with 1.7% DMSO. After 4 weeks in culture, differentiated HepaRG cells were seeded at a high density of approximately 227’000 cells/cm^2^ (72,000 cells/well with 100 μl per well) into black 96-well plates with a clear bottom (cat.nr.: 3603, Corning, New York, USA) and maintained for 10 days. After 10 days, cells were first adapted to treatment medium (culture medium containing 0.5% DMSO and 2% FBS) for 48 h and then used for cell viability and high-content imaging analysis.

### Cell viability assessment

Cell viability was analyzed using the WST-1 assay (cat. nr.: ab155902, Abcam, Cambridge, UK). HepaRG cells were exposed to chemicals at defined concentrations (**Table 2**) for 24 h. Triton X-100 (0.01%) was a positive control. One hour before the end of incubation period, 10 μl of WST-1 reagent was added to each well, quickly mixed by shaking the plate, and returned to the cell culture incubator (37 °C, 5% CO_2_). After 1 h, absorbance was measured (450 nm, reference wavelength 620 nm) using a plate reader (Infinite M200 Pro, Tecan group, Männerdorf, Switzerland). Absorbance was then subtracted from the reference wavelength, corrected for background by subtracting values measured without cells, and then normalized to the solvent control, which was set to 100 %. Dose-response data was fitted using drc package (Version 3.0.1) (Ritz *et al*., 2015) in R 3.6.1. Data are from at least 3 independent experiments with 6 technical replicates per condition.

### High-content imaging assay

Cellular responses were quantified using a high-content imaging approach. Based on cell viability assessment, concentrations below the EC_50_, used as a reference for overt toxicity, were tested in the high-content imaging assay. After chemical exposure, live cells were incubated with 5 μM CellROX (cat. nr.: C10422, ThermoFisher Scientific, Massachusetts, USA) by adding 10 μl of a 55 μM CellROX solution in treatment medium directly to the well. After incubation of 30 min at 37 °C, 5% CO_2_, cells were rinsed twice with treatment medium without serum. After removal of the treatment medium, cells were incubated with 50 nM Mitotracker Orange CM-H2TMROS (cat. nr.: M7511, ThermoFisher Scientific, Massachusetts, USA) for 10 minutes at 37 °C, 5% CO_2_. Subsequently, they were washed 3 times with PBS and fixed with 4% PFA at room temperature for 20 min. After washing 3 × 5 minutes, cells were stained with 50 nM Bodipy (cat. nr.: D3921, ThermoFisher Scientific, Massachusetts, USA) and 2 μM Hoechst (cat. nr.: 62249, ThermoFisher Scientific, Massachusetts, USA) at room temperature for 15 min, rinsed once with PBS (cat. nr.: 14190250, ThermoFisher Scientific, Massachusetts, USA), and then kept in PBS. Images were acquired using an ImageXpress micro (IXM) from Molecular Devices with a 20x 0.75 NA S Fluor objective, and 25 images were captured per well. The photometrics CoolSNAP HQ camera was set to 16 bit with a binning of one and a pixel size of 6.45 μm x 6.45 μm.

### Automated quantitative image analysis

Images were quantified using the open source software CellProfiler 3.1.8 (McQuin *et al*., 2018) and ImageJ 1.52g (Schindelin *et al*., 2012). Image analysis pipelines used in this publication are available as supplementary materials and are briefly described here. For cell population level data, nuclei were segmented using Otsu thresholding (Sankur, 2004) and everything outside the diameter range of 10-100 pixel was discarded. For lipid droplet quantification, GFP images were first deconvoluted using ImageJ 1.52g and then imported into the CellProfiler pipeline. A binary mask of the lipid droplets then was created, segmented, and quantified using the standard settings in the “IdentifyPrimaryObjects” module (**Supplementary figure 1**). Next, mitotracker and CellROX image intensity were measured. All parameters were exported to Excel spreadsheets and analyzed in R 3.6.1, as further explained in the paragraph “Dose response data and model fitting” below. For the single cell analysis pipeline, nuclei images were used to identify individual cells. Cell borders were determined based on the mitotracker staining with the propagation method (Jones *et al*., 2005). All quantifications from the cell population level pipelines were also performed at the single cell level. In addition, the following parameters were quantified per cell: size and shape of identified objects, object intensity, and object textures. The data was then exported to an SQLite database and analyzed using R 3.6.1. These pipelines can be used for a rapid population-based quantification which is amenable for high throughput assay or for a detailed single cell analysis to identify heterogenous responses and distinct cellular phenotypes.

### Data analysis of single cell data

The single cell SQLite database file was read using RSQLite p (Kirill *et al*., 2020). Data across three independent experiments with 6 technical replicates per concentration were summarized in a data matrix. This data matrix consisted of over 10^6^ cells with over 800 features per cell. It was imported into R 3.6.1. Constant and NA values were removed from the data matrix in order to perform a principle component analysis. Subsequently, the parameter cytoplasm area size was used to separate hepatocyte-like cells (<5000 px^2^) and cholangiocyte-like cells (>5000 px^2^) with the assumption that cells with a cytoplasm area below 5000 px^2^ are likely to be hepatocytes and those above likely to be cholangiocytes. Separation by size was further validated by anti-ASGPR1 immunostaining, a basolaterally expressed surface protein highly specific to hepatocytes (**Supplementary figure 2**). Its main function is the clearance of desialylated glycoproteins (D’Souza and Devarajan, 2015). Principle component analysis was used for dimensionality reduction and performed on the hepatocyte data set and plotted using ggplot2.

### Machine learning approach to quantify apoptotic cells

Apoptotic cells were quantified using a supervised machine learning approach. Nuclei images were analyzed with CellProfiler and intensity, size, shape and texture features were extracted. These data were loaded into CellProfiler Analyst (Jones *et al*., 2008) together with the raw nuclei images. A random forest classifier was trained to distinguish between healthy and apoptotic cells using a training set consisting of 477 different nuclei (**Supplementary figure 3A**). To take imbalance in the dataset into account, random undersampling was performed in the training set to achieve a 67/33 ratio (323 healthy, 154 apoptotic). The training set was assembled using nuclei from the menadione exposure across three independent experiments. Apoptotic nuclei were defined based on morphological nuclear fragmentation. The model was trained until a classification accuracy between 80 and 90 % was reached. Classification accuracy was determined based on user-defined assessment. In addition, images from the positive control were visually inspected to check the overall classification performance of the model and were corrected if numerous misclassifications were identified. If the classification was satisfying, the entire dataset was then scored. The resulting hit table was imported into R and the number of apoptotic cells were normalized to total cell number in the same well and further to untreated cells (**Supplementary figure 3**). Image analysis and supervised machine learning were performed on a terminal server based on Windows 10 equipped with 40 cores Intel Xeon Gold 6150 @ 2.7 Ghz and 512 GB RAM.

### Dose-response data and model fitting

Dose-response data were collected on a population level (i.e., per well averages) and combined with responses from all HepaRG cells per well. For every biological endpoint, the following raw parameters were selected per well and normalized to cell number per well:

- Lipid accumulation: Total number of lipid droplets divided by total number of cells
- Mitochondrial membrane potential: (Mean intensity–background intensity)/total number of cells
- Oxidative stress: (Mean intensity–background intensity)/total number of cells
- Nuclear morphology/cell death: number of apoptotic cells/total number of cells

These data were normalized to untreated cells to derive a fold change; unexposed cells had a fold change of 1. All data were normalized per plate to overcome possible plate effects. Non-linear regression of dose-response data was performed by fitting a log-logistic model with 3 or 4 parameters. The best fitting model was selected based on lowest Akaike Information Criterion (AIC) (Akaike, 1973) and visual inspection. A positive hit, i.e., an induction of a biological endpoint by exposure, was identified when the non-linear regression exceeded the calculated threshold of ± 2 standard deviations of the response at the two lowest concentrations. Dose-response data were fitted using the drc package (Ritz *et al*., 2015) and plotted with ggplot2 (Hadley Wickham, 2016). The R script of dose-response data and model fitting is available as supplementary material.

### Benchmark concentration modeling

In order to determine a point of departure of the high-content imaging concentration-response data, benchmark concentration (BMC) was modelled and calculated using the benchmark dose software (BMDS) version 3.1.2 (Gift *et al*., 2019). The benchmark dose technical guidance was used as a basis to perform the analysis (John *et al*., 2012). Normalized data including tested concentrations were entered as individual data points into the BMDS Excel application. The BMC was then defined as a benchmark response (BMR) of a change in the mean of one standard deviation from the control mean. Concentration-response data were then fitted to several models including exponential, hill, linear, polynomial and power models. Normal and log-normal distribution with constant or non-constant variance were also tested to find the best fit. BMC and its corresponding 95% confidence interval (95% CI) were then calculated. Selection criteria for the model with the best fit were the Akaike Information Criterion (AIC), goodness of fit *p*-value, scaled residuals for dose group near BMD, and for control dose group and BMDS recommendation. If there were several models which received a BMDS recommendation, BMCs from all models were averaged. BMCs were calculated for all chemicals tested (**Table 3**).

**Table 3.** Comparison of benchmark concentration and predicted Css with published pharmacokinetic values. Where indicated, in vivo derived Css values were calculated using following formula: Css = (Dose*F)/(Cl*dosing interval) or Css = k0/Cl with F = bioavailability, Cl = clearance, k0 = infusion rate and Css = steady state concentration.

| Chemical | LA<sup>1</sup> | MMP<sup>1</sup> | OS<sup>1</sup> | NM<sup>1</sup> | Median C<sub>ss</sub><sup>2</sup><br>(μM) | *In vivo* C<sub>ss</sub> (μM) | Species | *In vivo* C<sub>ss</sub><br>reference |
| --- | --- | --- | --- | --- | --- | --- | --- | --- |
| lomitapide | - | 0.58 | 10.5 | 4 | 0.0017 | 0.0002<sup>3</sup> | human | (Aegerion Pharmaceuticals, 2012) |
| fialuridine | - | 6.3 | - | 45.1 | 0.2 | 0.6<sup>3</sup> | human | (Bowsher *et al.*, 1994) |
| menadione | 2.8 | - | - | 8.1 | 5.7 | 12.6<sup>4</sup> | rabbit | (Hu *et al.*, 1996) |
| caffeine | - | - | - | - | 7.8 | 10.8 – 53.6 | human | (Birkett and Miners, 1991) |
| WY-14643 | 40.98 | 38.1 | - | 386.3 | 66 | 2.5<sup>3,5</sup> | rat | (Cunningham, 2007) |
| metformin | 987.1 | 16'887 | - | 1936.5 | 9.6 | 5.2 | human | (Graham *et al.*, 2011) |
| β-naphthoflavone | - | - | - | - | 32 | 2.3<sup>3</sup> | rat | (Adedayo Adedoyin, Leon Arons, 1993) |
| amiodarone | 8.5 | 13.8 | 5.35 | 14.95 | 5.2 | 5.4 – 6.0 | human | (Freedman and Somberg, 1991) |
| tetracycline | 32.9 | 218.43 | 69.7 | 32.7 | 12.6 | 9.7<sup>3</sup> | human | (Welling *et al.*, 1977) |
| valproic acid | 569.95 | 1311.3 | 1379.8 | 611.6 | 203.1 | 346.7 – 693.4 | human | (Vasudev *et al.*, 2001) |

LA: lipid accumulation, MMP: mitochondrial membrane potential, OS: oxidative stress, NM: nuclear morphology, <sup>1</sup>Benchmark concentration (μM) per imaging endpoint calculated with BMDS software, <sup>2</sup>Derived from IVIVE, <sup>3</sup>calculated with formula described above, <sup>4</sup>in rabbits, <sup>5</sup>in rats

### *In vitro* to *in vivo* extrapolation (IVIVE)

IVIVE was performed based on the previously published method by Rotroff and Wetmore et al. (Rotroff *et al*., 2010; Wetmore *et al*., 2015) and using the high-throughput toxicokinetic (httk) R package (version 1.10.1) developed by the US-EPA. The httk package contains four toxicokinetic models which can be parameterized using high throughput-derived *in vitro* data on plasma protein binding and hepatic clearance. Moreover, it has a Monte Carlo sampler to simulate population variability and includes tools for reverse dosimetry together with functions for the analysis of concentration versus time simulations (Pearce *et al*., 2017). In our study, a three-compartment steady state pharmacokinetic (PK) model was parameterized and used for the simulations. For all simulations, the following assumptions were made: oral route of exposure, daily dosing with constant dose rate, and 100 % bioavailability. Human data was used except where otherwise stated. Steady-state concentration (C_ss_) in the blood was calculated using formula 1 (Pearce *et al*., 2017)

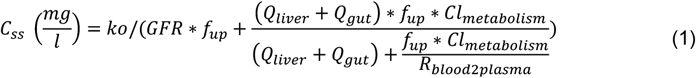

where ko = constant dose rate (mg/kg BW/day), f_up_ = *in vitro* measured chemical fraction unbound in plasma, Cl_metabolism_ = metabolic clearance scaled from *in vitro* intrinsic hepatic clearance, R_blood2plasma_ = ratio of blood concentration of a chemical to the plasma concentration, GFR = glomerular filtration rate (mean 5.17 ml/min/kg^3/4), Q_liver_ = blood flow to the liver (mean 59.9 ml/min/kg^3/4), and Q_gut_ = blood flow to the gut (mean 47.5 ml/min/kg^3/4). Fraction unbound, metabolic clearance, and R_blood2plasma_ values were obtained from the httk package or from literature. Recognizing that formula 1 is linear in dose rate (ko), oral equivalent doses (OED) were then calculated using the C_ss_ predicted for a 1 mg/kg/day dose rate as in formula 2 (Rotroff *et al*., 2010).

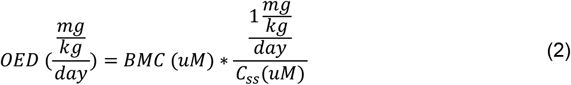

The calculated OED corresponds to an oral daily dose which would lead to a plasma concentration equal to the *in* vitro-derived BMC. The OED is linearly related to the BMC and inversely related to C_ss_. This equation is only valid for first-order metabolism (Rotroff *et al*., 2010). A Monte Carlo analysis was used to simulate population variability in, GFR, Q_liver_, Q_gut_, and Cl_metabolism_ across 1000 healthy individuals. Extrapolations were performed for each chemical and endpoint which was positive in the imaging assay. Every predicted parameter per chemical was then combined into one boxplot per endpoint, which contained all predicted OEDs across all positive endpoints.

For the model evaluation, C_ss_ of selected reference chemicals were predicted using formula 1 with the same dose rate from the *in vivo* C_ss_ measurements. For chemicals with no in vitro intrinsic hepatic clearance and fraction unbound values available in the database included in the httk package, in silico predictions from (Sipes *et al*., 2017) were loaded using load_sipes2017(). In addition to that, in vitro intrinsic hepatic clearance values were calculated for a selection of chemicals (Lomitapide, Fialuridine, Metformin, beta-naphthoflavone and Menadione.) using *in vivo* clearance values (**supplementary table 1**) and formula 3:

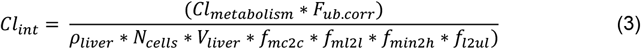

where Cl_int_ = in vitro intrinsic hepatic clearance in ul/min/10^6 cells, Cl_metabolism_ = *in vivo* clearance in L/h/kg (if available hepatic otherwise total clearance), F_ub.corr_ = assay correction factor which is assumed to be 1 (i.e., no corrections), ρ_liver_ = liver density of 1.05 g/ml, N_cells_ = 1.1*10^8 hepatocytes per gram of liver, V_liver_ = liver volume: 0.0245 L/kg (human), 0.0349 L/kg (rat), 0.04 L/kg (rabbit), f_mc2c_ = conversion factor for millions of cell to one cell, f_ml2l_ = conversion factor from ml to l, f_min2h_ = conversion factor from minute to hour, and fl2ul = conversion factor from L to ul. The formula to calculate Cl_int_ was derived from the function calc_hepatic_clearance () within the httk package. Predicted C_ss_ values were then compared to published *in vivo* C_ss_ values. If there was no published *in vivo* C_ss_ available, C_ss_ was then calculated based on published *in vivo* pharmacokinetic parameters using the following formula (**Table 3**):

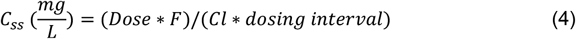

Vice versa, predicted OED derived from BMC were compared to published exposure values of the corresponding chemicals. Simulations were performed in humans where pharmacokinetic parameters for humans were available; otherwise, estimations were calculated in rats or rabbits, as stated in figure 7. The R script written to perform the IVIVE analysis is available as supplementary material.

## 4 Results

### Four hallmarks of steatosis were simultaneously quantified by high-content imaging

A high-content imaging assay was developed for simultaneous quantification of four molecular hallmarks: lipid accumulation, mitochondrial membrane potential, oxidative stress, and nuclear morphology as a proxy for cell death. HepaRG cells were selected as a biological model due to their unlimited growth capacity and phenotypic stability, as well as the stability of enzymatic activities over 4 weeks of cultivation, thus having similar metabolic function to primary human hepatocytes, but overcoming limitations such as donor variability and phenotypic instability associated with the use of primary cells (Andersson *et al*., 2012; Aninat *et al*., 2006; Lambert *et al*., 2009; Lübberstedt *et al*., 2011; Guillouzo *et al*., 2007). The model contains two cell types, hepatocyte-like and cholangiocyte-like cells (Parent *et al*., 2004), which allows elucidation of cell-specific mechanisms of toxicity (Jossé *et al*., 2008). Thus, differentiated HepaRG cells were exposed to test chemicals and live cells were stained for oxidative stress and mitochondrial membrane potential using CellROX deep red and Mitotracker Orange, respectively. CellROX deep red fluoresces upon oxidation. Oxidative stress was then measured on the basis of CellROX intensity, which reflects hydroxyl radical and superoxide anion levels (ThermoFisher Scientific). Mitotracker Orange CM-H2TMROS accumulation depends on mitochondrial membrane potential and fluoresces upon oxidation. Both dyes were retained after aldehyde fixation. Mitochondrial membrane potential was quantified by measuring the intensity of the Mitotracker dye, and a decreased intensity was diagnostic of decreased mitochondrial potential. Finally, cells were fixed and stained for lipid droplets with Bodipy 505/515 and nuclei with Hoechst, which was then used to count the number of cells analyzed. An automated image analysis platform was developed to quantify the four endpoints on a population-based level and machine learning was implemented to use nuclear morphology as a proxy to quantify cell death. Single cell level analysis was used to investigate different response patterns in hepatocytes and cholangiocyte-like cells upon chemical exposure.

### Multiparametric analysis of reference chemicals

To validate the assay, positive and negative control chemicals with previously established mechanisms of action were characterized by the high content imaging approach described above (**Figure 1A**). Thus, we tested a mixture of oleic acid and palmitic acid, which are often used to induce steatosis in vitro (Sharma *et al*., 2011; Graffmann *et al*., 2016; Gómez-Lechón *et al*., 2007; Michaut *et al*., 2016); rotenone, a selective complex I inhibitor known to decrease mitochondrial membrane potential (Siddiqui *et al*., 2013) and therefore used as a positive control for inducing mitochondrial dysfunction via a decrease in mitochondrial membrane potential; amiodarone, which induces oxidative stress (Anthérieu *et al*., 2011; Tolosa *et al*., 2016); menadione, which induces DNA damage and cell death (Loor *et al*., 2010); and caffeine as negative control (Persson *et al*., 2013; Saito *et al*., 2016). Concentration ranges were selected on the basis of preliminary cell viability assessment, and each chemical was used at concentrations ranging up to the corresponding EC_50_ value (**Supplementary figure 4**). Cell responses were evaluated on the basis of an automated image acquisition workflow and automated image analysis was then used to quantify the cellular responses of the four endpoints (**Figure 1B**). To evaluate responses on a single cell basis, over 800 different features were extracted from every single cell. The resulting multidimensional dataset was then analyzed using principal component analysis for dimensionality reduction (**Figure 1B**). For endpoints that exceeded a calculated threshold of ±two times the standard deviations of the response to the two lowest concentrations were further evaluated by IVIVE.

**Figure 1.**
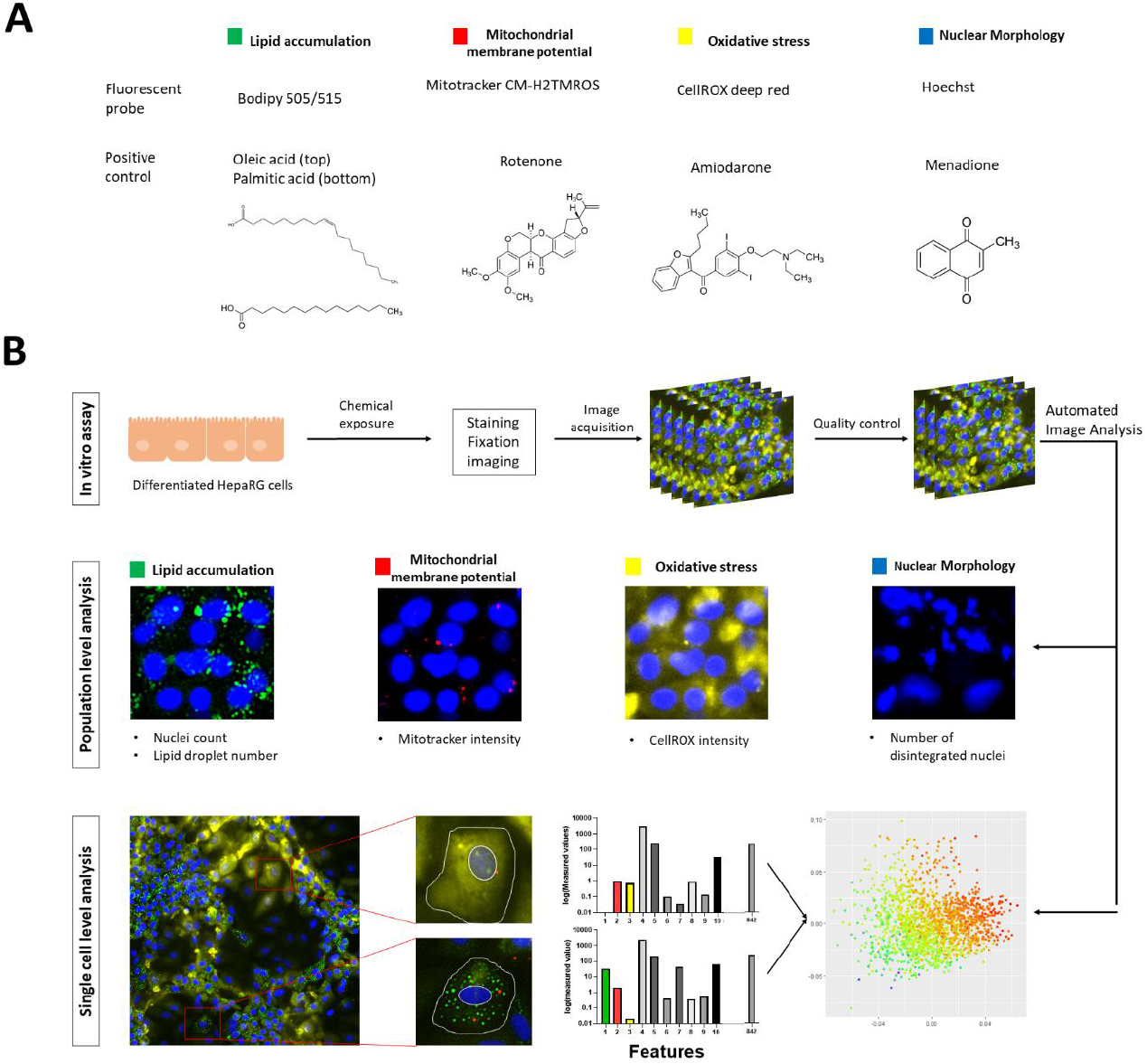
Basis of high-content imaging assay. A) Fluorescent probes for the selected endpoints including corresponding positive controls. B) Description of the workflow including rapid population-based quantification and detailed single cell analysis. During single cell level analysis, over 800 different features (intensity, size/shape, and texture) per cell were quantified and analyzed using principal component analysis (PCA). Plotted data is then color labelled by using the values from following parameters: lipid droplet size, mitotracker intensity and CellROX intensity.

**Figure 2.**
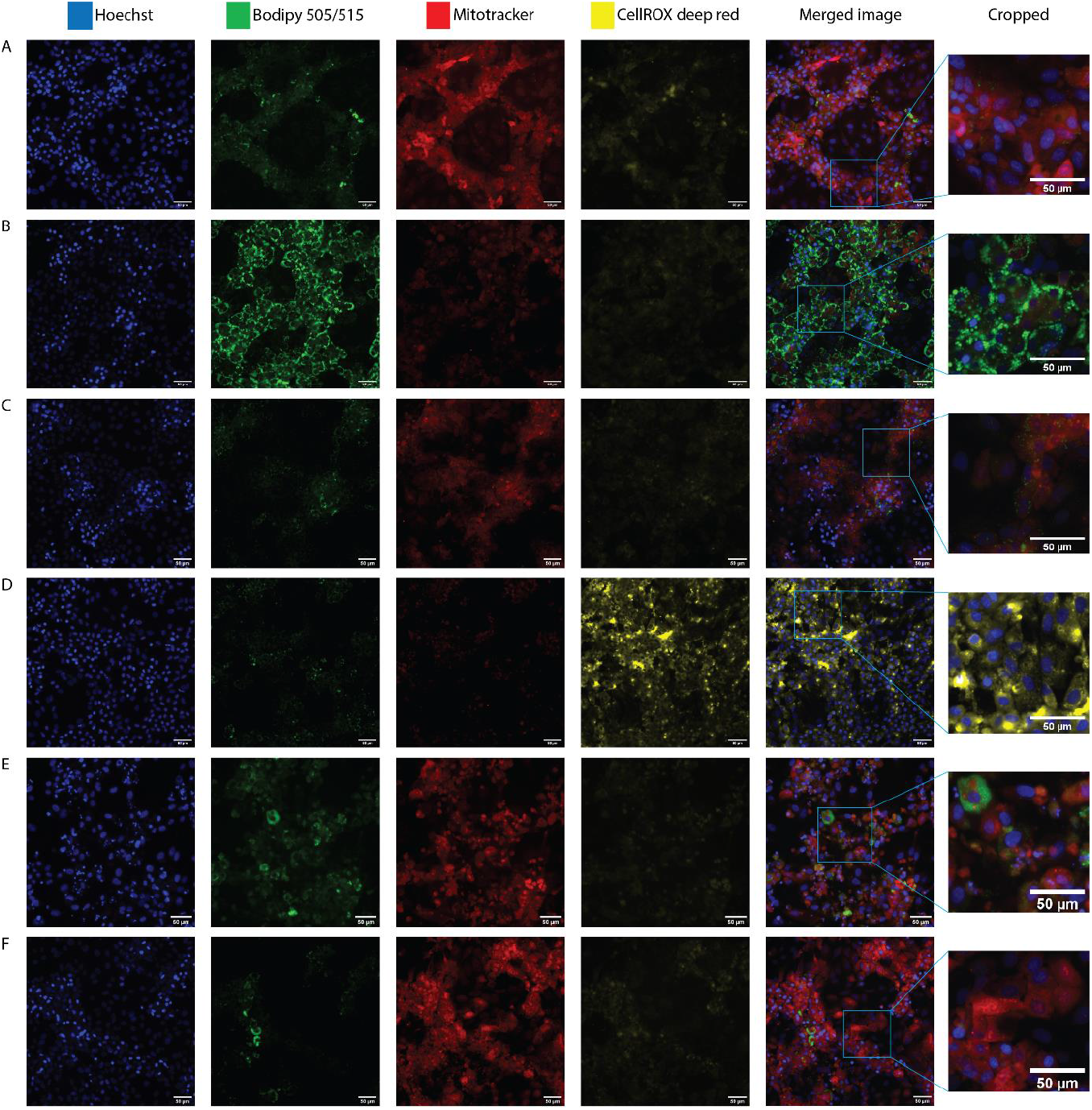
Representative fluorescent images from chemically exposed HepaRG cells. A) DMSO control B) Oleic acid/palmitic acid (1000 μM) C) Rotenone (20 μM) D) Amiodarone (50 μM) E) Menadione (50 μM) F) Caffeine (5mM). Uncropped images were acquired at 20x. Scalebar = 50 μm. Blue=nuclei, green=lipid accumulation, red=mitochondrial membrane potential, yellow=oxidative stress.

Response profiles of the test compounds were consistent with expected outcomes. Lipid accumulation increased in a dose-dependent manner upon exposure to all four positive control chemicals. The most potent exposure was oleic acid/palmitic acid (1000 μM), which induced a 2.3-fold-change in lipid accumulation. Caffeine induced a decrease of lipid accumulation that exceeded the threshold only at the highest concentration (5000 μM). In the case of mitochondrial membrane potential, oleic acid/palmitic acid, rotenone, and amiodarone all induced a dose-dependent decreasing response which exceeded the threshold at 62.5, 25 and 25 μM respectively, whereas menadione and caffeine resulted in no change in mitochondrial membrane potential at 24 h. In the case of oxidative stress, amiodarone exposure led to a dose-dependent increasing response and exceeded the threshold at 50 μM. In contrast, oleic acid/palmitic acid, rotenone, menadione, and caffeine did not alter oxidative stress levels. The number of apoptotic cells (nuclear morphology) increased in oleic acid/palmitic acid-, amiodarone- and menadione-treated HepaRG cells in a dose-dependent fashion, exceeding the threshold at 250, 25 and 25 μM. Neither rotenone nor caffeine exposure led to alterations in the number of apoptotic cells (**Figure 3**). In previous studies, HepaRG cells were found to report accurately on chemical-induced steatosis, oxidative stress, and mitochondrial dysfunction (Allard *et al*., 2020; Anthérieu *et al*., 2011; Angrish *et al*., 2017; Tolosa *et al*., 2016), and the positive and negative control chemicals tested here showed similar *in vitro* response patterns as previously described (Sharma *et al*., 2011; Graffmann *et al*., 2016; Siddiqui *et al*., 2013; Anthérieu *et al*., 2011; Gómez-Lechón *et al*., 2007; Michaut *et al*., 2016; Tolosa *et al*., 2016). These data confirmed the accuracy of the in vitro assay and data analysis pipelines.

**Figure 3.**
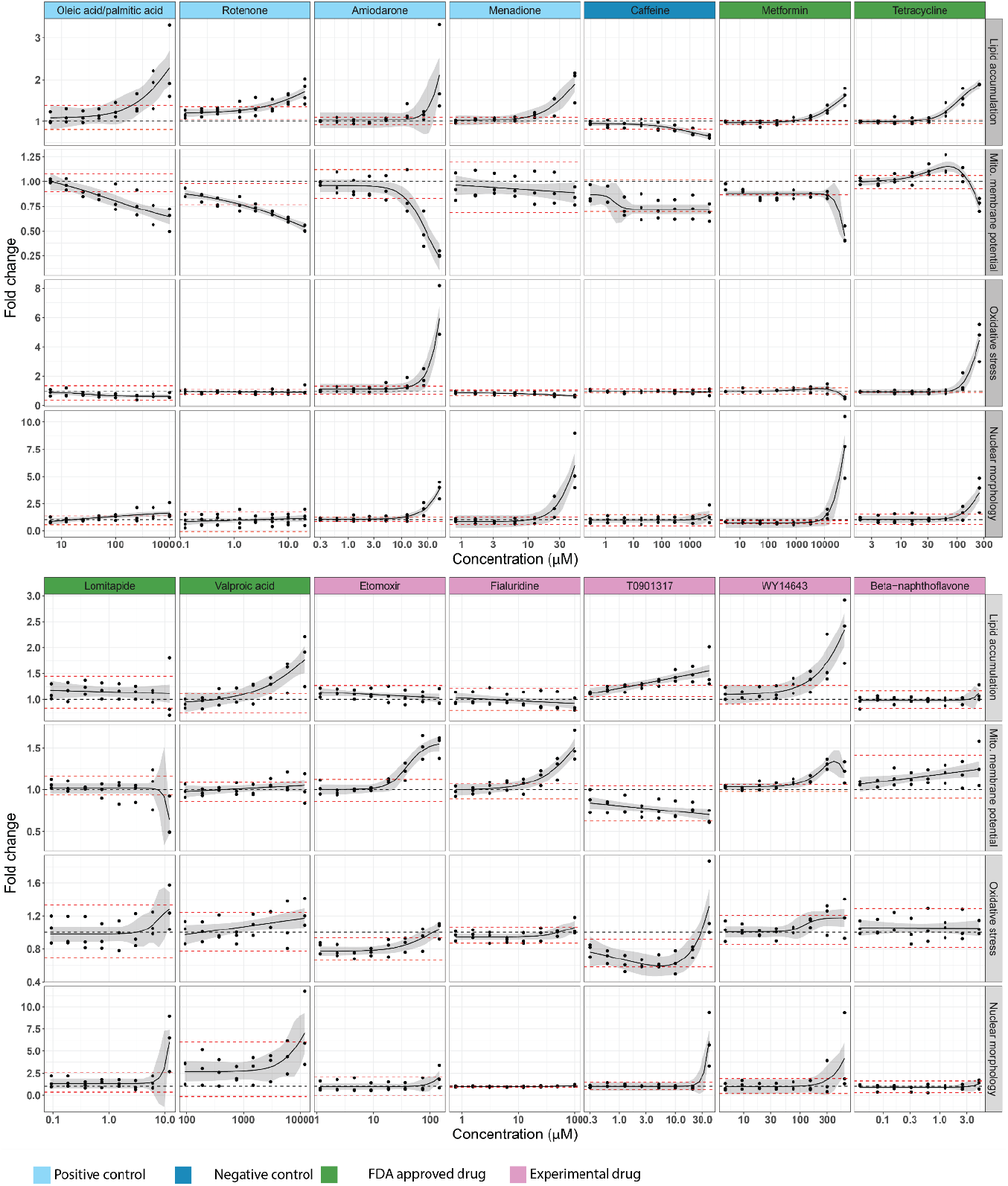
Cellular responses for four key steatosis-relevant endpoints. Grey area around non-linear regression represents 95% CI. Black dotted line represents a fold change of 1 (DMSO control). Red dotted lines represent threshold (± 2 standard deviations of the response at the two lowest concentrations). All data were first normalized to cell number and then to untreated cells. At least 3 independent experiments with 6 technical replicates per concentration per experiment were conducted for all chemicals.

Following the effective systematic characterization of positive and negative control chemicals for each endpoint, we tested four Food and Drug Administration (FDA)-approved drugs with known mechanisms of action. They included metformin, an anti-diabetic drug which inhibits mitochondrial complex I (Pernicova and Korbonits, 2014), tetracycline, an antibiotic drug that decreases β-oxidation, inhibits microsomal triglyceride transfer protein (MTP) and upregulates PPARγ and SREBP1-c (Schumacher and Guo, 2015; Anthérieu *et al*., 2011), lomitapide, an anti-hypercholesterolemic drug and inhibits MTP (Lin *et al*., 2014), and valproic acid, an anti-convulsant drug that inhibits β-oxidation and upregulates CD36 and DGAT2 (Schumacher and Guo, 2015; Bai *et al*., 2017). Lipid accumulation increased in a dose-dependent manner above the threshold after exposure to metformin, tetracycline and valproic acid, whereas no effect was observed for lomitapide. Mitochondrial membrane potential decreased when cells were exposed to metformin or lomitapide, but only at the highest concentrations (50,000 and 12 μM respectively). An unexpected hormetic effect was observed in cells exposed to tetracycline; between 31.25 and 125 μM, a hyperpolarization, which to our knowledge has not been reported previously, was observed and only at the highest concentration (250 μM) did the mitochondrial membrane potential decrease. No changes in MMP were measured in valproic acid-exposed cells. Oxidative stress dose-dependently increased after tetracycline exposure and exceeded the threshold at 125 and 250 μM. A decrease of oxidative stress below the threshold was observed at 50’000 μM metformin. There were no observed alterations in oxidative stress levels for lomitapide- and valproic acid-exposed cells. Finally, the number of apoptotic cells increased above the threshold in metformin, tetracycline and lomitapide treated cells (**Figure 3**). These data thus confirm the anticipated activation profiles of the cellular markers profiled in parallel for FDA-approved drugs with known mechanisms of action.

As a next step, we were interested to test etomoxir and fialuridine, both chemicals that had their clinical development terminated due to severe hepatotoxicity in clinical trials, but with limited knowledge of the underlying mechanisms (Holubarsch *et al*., 2007; Manning and Swartz, 1995). Etomoxir was developed as an anti-diabetic drug and is an irreversible carnitine palmitoyltransferase 1 (CPT1) inhibitor (Merrill *et al*., 2002). Fialuridine is a nucleoside analogue that was developed to treat chronic hepatitis and inhibits mitochondrial DNA polymerase γ (Lewis *et al*., 2003; McKenzie *et al*., 1995). No lipid accumulation was observed in etomoxir-nor fialuridine-exposed cells. MMP increased in fialuridine and etomoxir exposed cells and exceeded the threshold at 25 and 36 μM respectively. Moreover, oxidative stress levels in etomoxir-exposed cells were lower than baseline at low drug concentrations (1.15 – 18.1 μM) and increased to baseline at higher drug concentrations (36.25, 72.5 and 145 μM), whereas fialuridine did not induce oxidative stress above the threshold. Both chemicals also did not increase the number of apoptotic cells (**Figure 3**).

As a step to characterize the in vitro assay, we were interested in testing chemicals known to bind to nuclear receptors, as nuclear receptor activation is the molecular initiating event of steatosis. Thus, we characterized three experimental drugs: T0901317 is a synthetic liver X receptor agonist that increases the expression of SREBP-1c and ChREBP (Mitro *et al*., 2007; Cha and Repa, 2007), as well as WY14643, a dual PPARα/PPARγ agonist that increases oxidative stress in mouse liver and affects fatty acid metabolism (Jennings *et al*., 2014; Woods *et al*., 2007), and β-naphthoflavone (BNF), an aryl hydrocarbon receptor agonist that upregulates CD36 (Jennings *et al*., 2014; Lee *et al*., 2010). Lipid accumulation dose-dependently increased and exceeded the threshold for cells exposed to T0901317 and WY14643 but not BNF. Mitochondrial membrane potential dose-dependently decreased for cells exposed to T0901317, but did not exceed the threshold, whereas for WY14643, hyperpolarization of the mitochondrial membrane potential was observed at 150, 300, and 600 μM, whereas at the highest concentration (600 μM), an as-yet unprecedented hyperpolarization decrease was observed. Oxidative stress levels in T0901317-treated cells decreased at lower concentrations (0.3–20 μM) and were above the threshold only at the highest concentration (40 μM). No impact on oxidative stress within the tested concentration range was observed in cells exposed to WY14643 and BNF. The number of apoptotic cells increased when cells were exposed to T0901317 and WY14643 at the highest tested concentration (40 and 600 μM respectively), but not for BNF (**Figure 3**). Thus, the nuclear receptor agonists effectively induced lipid accumulation.

Of the 14 reference compounds tested, the outcomes were categorized as negative or positive, excluding amongst positive outcomes instances where no dose-response relationship was observed or if there was a cellular response only at the single highest dose tested, and whether that experimental outcome was expected vs. unexpected based on literature precedent

### Analysis of food-related chemicals and pesticides

Having validated the in vitro method for simultaneous quantification of steatosis-relevant cellular key events, we were interested in characterizing responses to food-related chemicals selected from structure-based nuclear binding prediction. To model receptor binding, about 6,000 Smiles were extracted from the regulated food-related use/occurrence chemical database from the Swiss Federal Food Safety and Veterinary Office and about 4,000 from the Toxcast database (Richard *et al*., 2016). We assessed the potential for nuclear receptor binding on the basis of molecular fragments and other relevant chemical features, by a previously reported workflow (Mellor *et al*., 2016), and identified about 80 chemicals positive for binding to one or more receptors. These 80 compounds were prioritized for testing on the basis of available *in vitro* and *in vivo* hepatotoxicity data and toxicokinetic data (**Supplementary table 3**). Uric acid, tartrazine, bisphenol A, atrazine, metazachlor, and vinclozolin gave rise to significant lipid accumulation, while for carbosulfan a decrease in lipid accumulation was observed at the highest tested concentration. Mitochondrial membrane potential increased in a dose-dependent manner in cells exposed to orotic acid, tartrazine, fructose and carbofuran, and exceeded the threshold for all except fructose. On the other hand, a dose-dependent decrease in mitochondrial membrane potential was observed in mepanipyrim-exposed cells but did not exceed the threshold. No response was observed in the number of apoptotic cells across all tested food-related chemicals and pesticides (**Figure 4**). These data emphasize the sensitivity of the cells for detecting the potential of chemicals to induce lipid accumulation and mitochondrial dysfunction without inducing cell death. Finally, although several chemicals contained structural alerts for receptor-binding, associated cellular responses were not observed, emphasizing the importance of experimental analysis of phenotypic endpoints.

**Figure 4.**
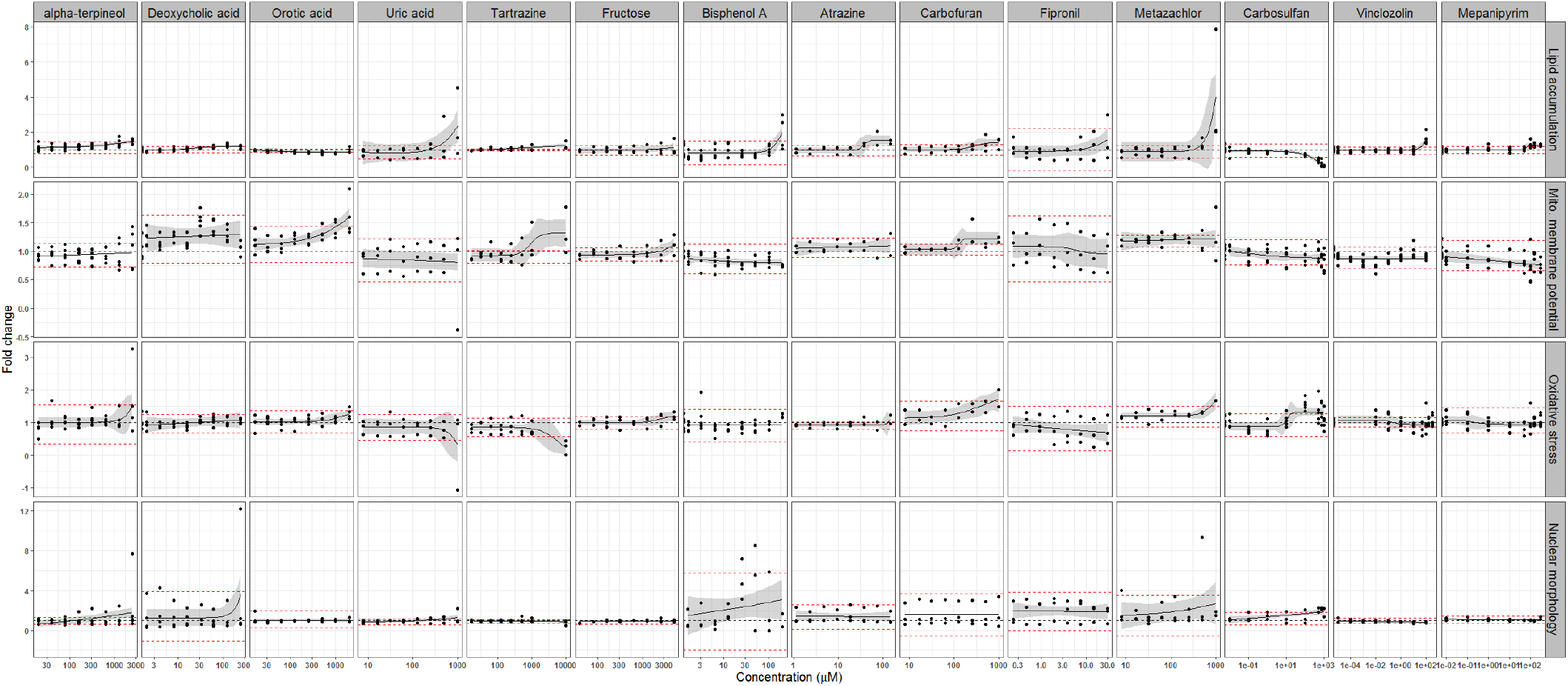
Cellular responses for four key steatosis-relevant endpoints of food-related chemicals. Grey area around non-linear regression represents 95% CI. Black dotted line represents a fold change of 1 (DMSO control). Red dotted lines represent threshold (± 2 standard deviations of the response at the two lowest concentrations). All data were first normalized to cell number and then to untreated cells. 2-3 independent experiments with 6 technical replicates per concentration per experiment were conducted for all chemicals.

### Chemical exposures induce distinct cellular endophenotypes within cell populations

On the basis of cell population-level responses, we could simultaneously quantify the four cellular responses, however, we could not distinguish specific responses of hepatocyte vs. cholangiocyte-like cells. Thus, to evaluate the potential heterogeneity of single cell responses and interrogate potential endophenotypes within and/or between cell populations in the HepaRG cells, we evaluated alterations of three of the key events (lipid accumulation, mitochondrial membrane potential and oxidative stress) in hepatocyte-like cells vs. cholangiocyte-like cells following tetracycline exposure. Tetracycline was selected because it induced distinct responses above the threshold in all four endpoints. We aimed to determine whether individual cell types accumulated higher numbers of droplets compared to controls, as well as how many cells had increased numbers of lipid droplets (**Figure 5**). In this way, we found that there were always more lipid droplets in hepatocyte-like cells compared to cholangiocyte-like cells at all tetracycline exposure concentrations, including background levels in unexposed cells (**Figure 5**). Interestingly, at 125 μM and 250 μM, the median of lipid droplets per hepatocyte-like cell did not drastically increase (~8 to ~12 lipid droplets per cell), but the number of hepatocyte-like cells with a larger median for droplet number than the median for droplet number in untreated cells increased. The opposite response was observed for cholangiocyte-like cells; namely upon exposure to 125 μM or 250 μM tetracycline, the median of lipid droplets per cholangiocyte-like cell increased, but the proportion of cholangiocyte-like cells with more lipid droplets remained unchanged (**Figure 5**). Similar to lipid droplets, the mitochondrial membrane potential was also elevated in hepatocytes compared to cholangiocyte-like cells for all tetracycline concentrations tested. Notably, at the highest concentration of tetracycline (250 μM), a decrease of mitochondrial membrane potential was observed in both cell types. Finally, for oxidative stress, similar patterns of changes were observed in hepatocytes which had increased levels of oxidative stress compared to cholangiocyte-like cells throughout all tested concentrations. At the highest concentration at which tetracycline was tested (250 μM), oxidative stress levels were elevated in both cell types. By evaluating increased oxidative stress on a single cell basis, it became apparent that the increased average values for oxidative stress on a cell population level arose from a small number of cells with extremely high levels of oxidative stress, rather than constitutive moderate increase across the whole cell population (**Figure 5**). In summary, hepatocytes tend to have an increased response compared to cholangiocyte-like cells and that rather a small population has a high response which drives population-based response levels.

**Figure 5.**
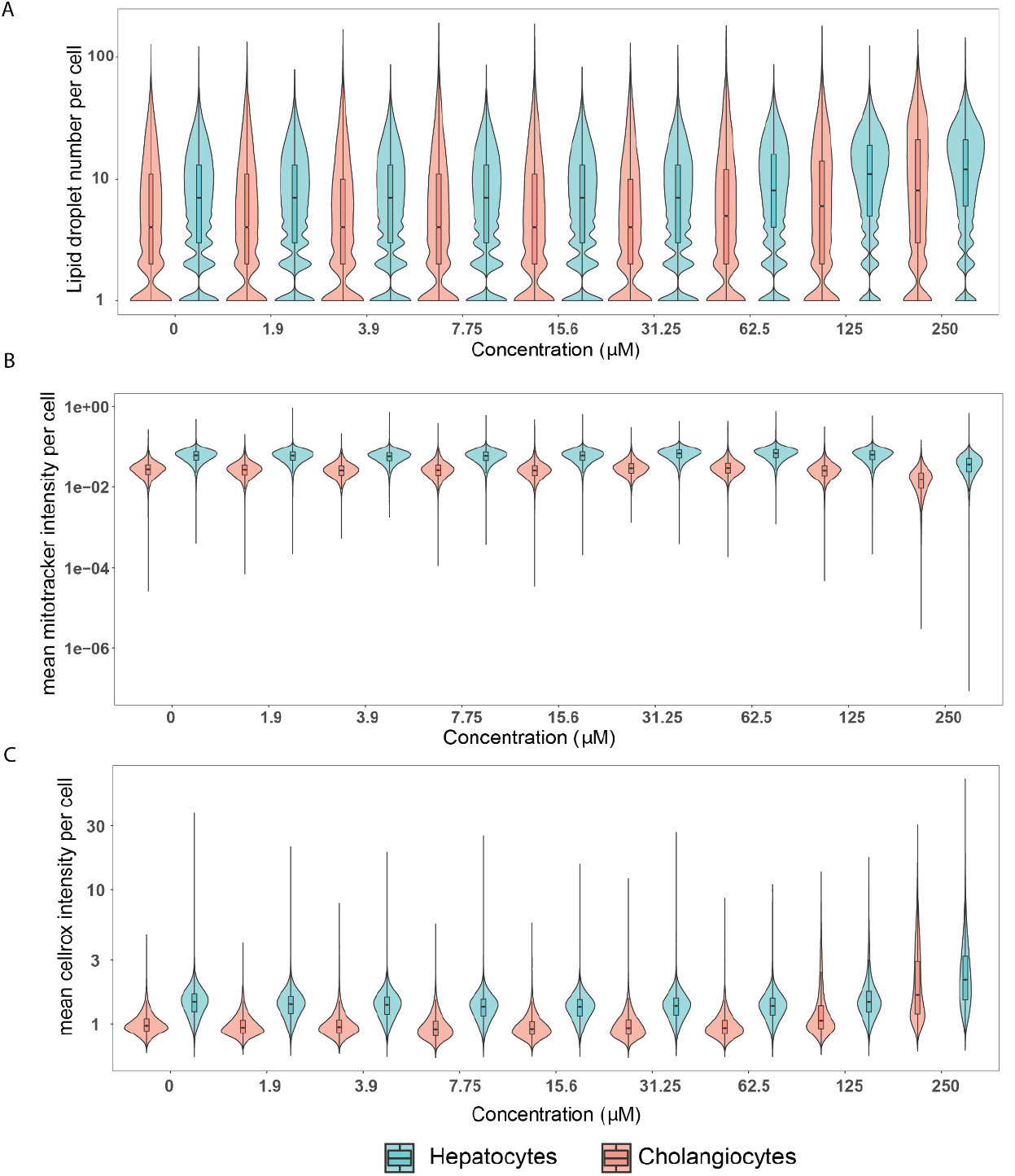
Quantification of selected endpoints in individual hepatocytes versus cholangiocyte-like cells after tetracycline exposure. The same raw data used in the population level analysis were also used for this single cell analysis. 30’000 to 150’000 cells per concentration and cell type were analyzed. Cells from 3 independent experiment were included in the analysis. Notched boxplots were overlaid in the violin plots to display the confidence interval around the median.

Given the heterogeneity of responses in hepatocytes and cholangiocyte-like cells, we investigated whether hepatocyte-like cells with a specific response in one endpoint (e.g., large lipid droplet size) responded similarly in other endpoints (mitochondrial membrane potential and oxidative stress) at the same concentration. For this purpose, over 800 features per cell were extracted by high content imaging and resulting PCA plots were color labelled with lipid droplet size, mitotracker intensity, and CellROX intensity to visualize their distribution (**Figure 6**). Heterogeneity in lipid droplet size and mitotracker intensity was observed in both chemically and mock-exposed hepatocyte-like cells, whereas it was only observed for oxidative stress in chemically exposed cells. Oxidative stress levels in non-exposed hepatocytes were similar across all measured hepatocytes (**Figure 6A**). While mitochondrial membrane potential did not decrease in hepatocytes with larger lipid droplets it did decrease in hepatocytes with smaller lipid droplets (**Figure 6B**). Similarly oxidative stress levels in chemically exposed hepatocytes with larger lipid droplets were lower than in hepatocytes with medium and small sized lipid droplets. Interestingly, a subgroup of hepatocytes with medium and small sized lipid droplets also had lower oxidative stress levels compared to other hepatocytes with the same lipid droplet size. Thus, tetracycline exposure led to heterogenous response patterns in hepatocytes involving a subpopulation that was protected from mitochondrial dysfunction but oxidative stress could be detected.

**Figure 6.**
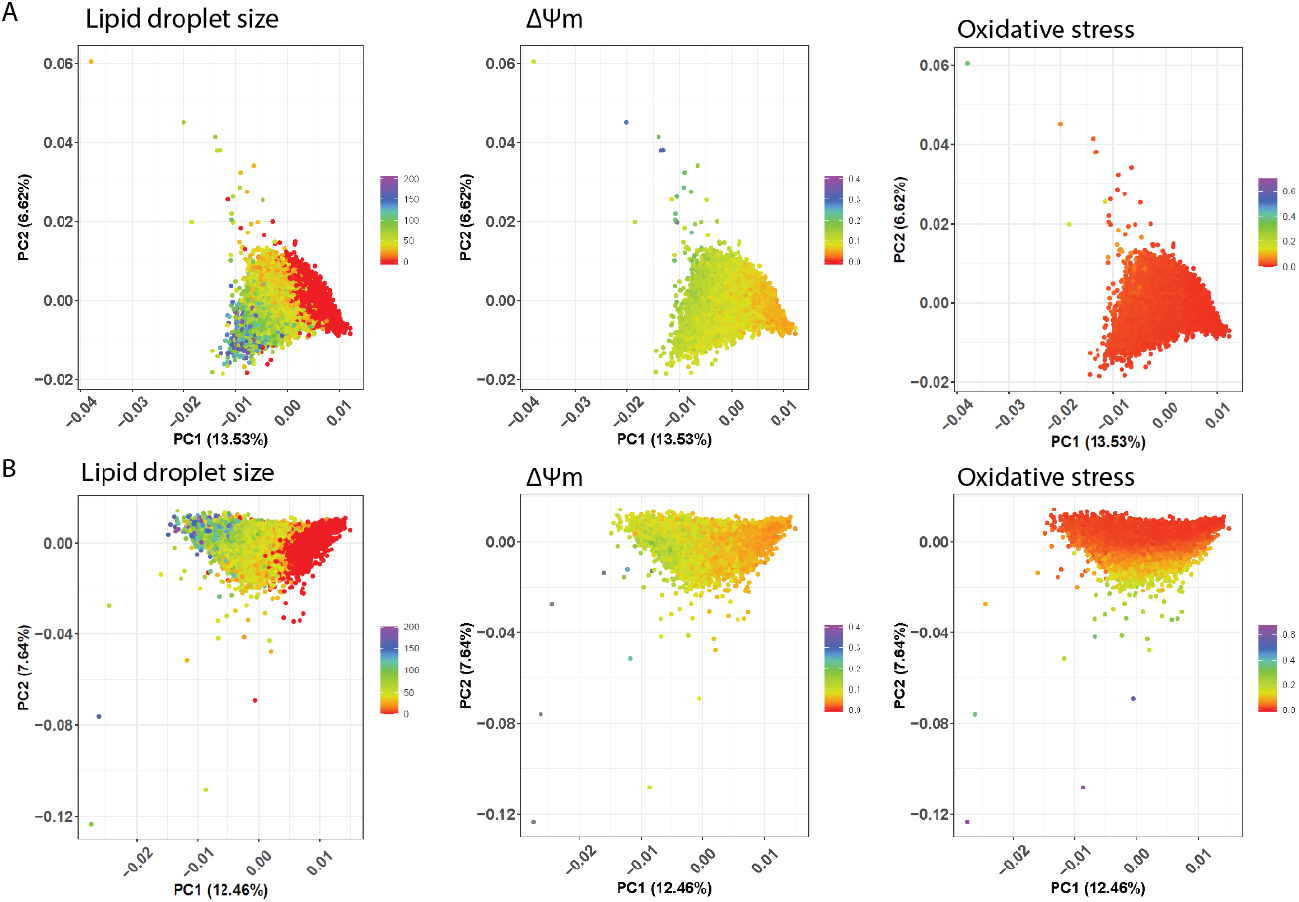
Heterogenous response pattern after tetracycline exposure in hepatocytes. A) untreated B) tetracycline (250 μM). A single point in the PCA plot represents a hepatocyte and its color code represents lipid droplet size (px^2), mitochondrial membrane potential (ΔΨm)(AU) or oxidative stress (AU). 80’000 to 125’000 cells from 3 independent experiments were analyzed per concentration.

### IVIVE

In order to evaluate the relevance of the *in vitro* concentrations that stimulated one or more of the endpoints, IVIVE was performed using pharmacokinetic modeling and reverse dosimetry. To gain an understanding of the model performance, predictions were made using *in vitro* PK data and a three-compartment steady state PK model for chemicals where *in vivo* PK data was available for evaluation. A Monte Carlo simulation modeling population variability across 1000 healthy individuals based on physiologies from a distribution of demographic and anthropometric quantities including sex, race/ethnicity, age, height and weight from the US Centers for Disease Control and Prevention National Health and Nutritional Examination Survey (NHANES) (Ring *et al*., 2017) yielded a range of plasma C_ss_ predictions consistent with a constant dose rate of 1 mg/kg bodyweight/day (**Figure 7A**). The predicted C_ss_ values were compared with *in vivo* C_ss_ (**Figure 7A**) which were derived from pharmacokinetic studies in human, rat or rabbit as indicated in (**Table 3**) For the ten chemicals, the coefficient of variation (R^2^) on a logarithmic scale was 0.78 and the root mean squared error was 0.69 (a factor of 4.9x). The chemicals with the greatest discrepancy where beta-naphthoflavone and WY-14643, although both *in vitro* and *in vivo* estimates of C_ss_ were in the μM range.

**Figure 7.**
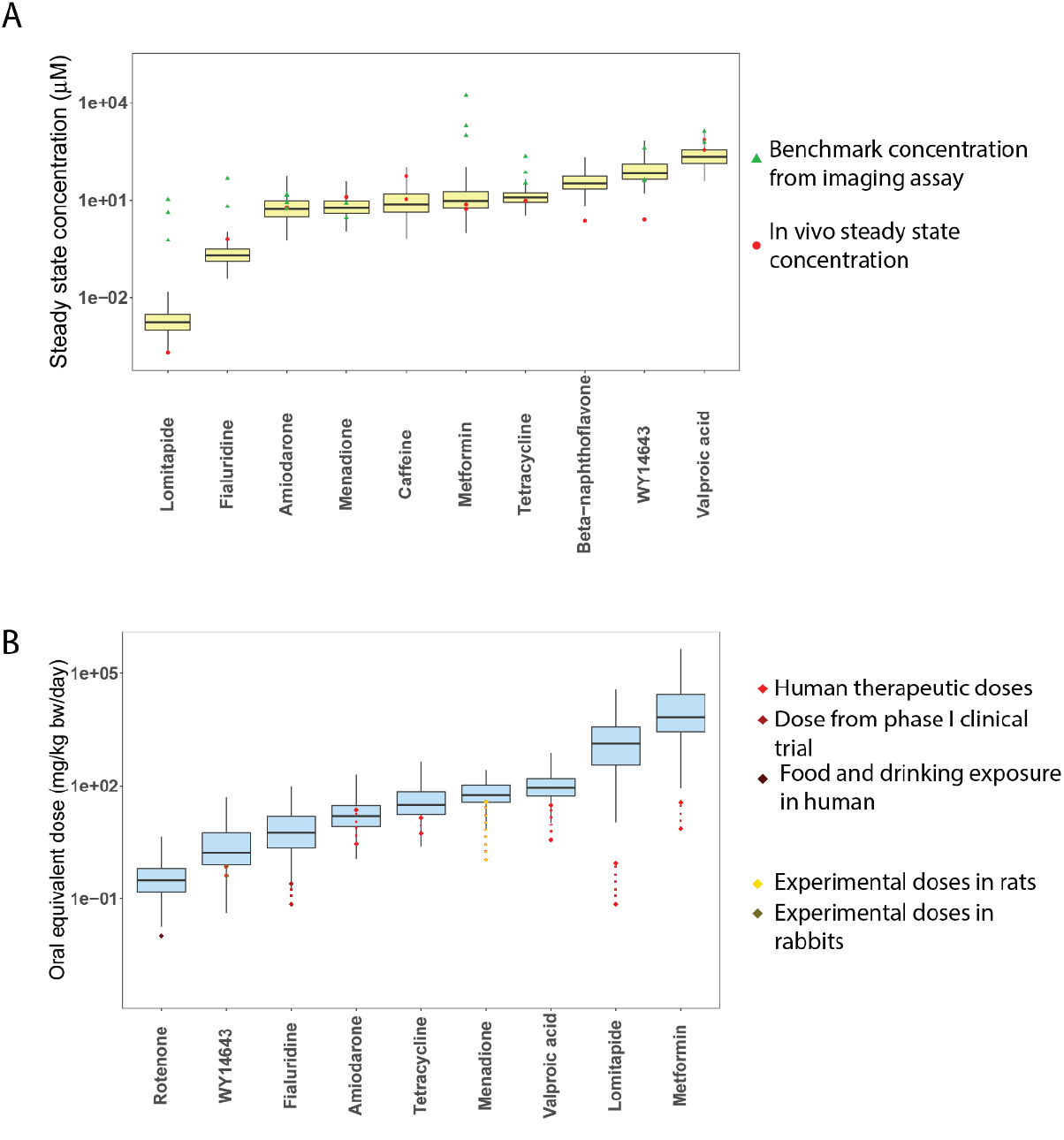
Pharmacokinetic modeling of selected reference chemicals. A) Forward dosimetry of chemicals with published Css values. Yellow boxplot represents predicted Css using IVIVE. B) Reverse dosimetry of chemicals with published human exposure values from different sources. Blue boxplot represents predicted oral equivalent doses using httk with a monte carlo population sampler. OED predictions were performed for every endpoint from the HCI assay which was above the calculated threshold.

Benchmark concentrations (BMC) were then calculated for every endpoint which was positive in the imaging assay from each chemical to determine a point of departure (**Table 3 and Table 5**). A direct comparison between the *in vitro* derived BMC and *in vivo* C_ss_ was then made to identify a potential *in vivo* hazard. For lomitapide, fialuridine, WY14643, metformin, and tetracycline, the calculated BMC was about 10 – 10’000 times higher than the median PBPK-predicted C_ss_. For amiodarone and valproic acid, the BMC was about 1.5-4 times higher than the median predicted C_ss_, whereas for menadione it was about 1.5-4 times lower than the median predicted C_ss_. For orotic acid, uric acid, carbofuran, and carbosulfan, the calculated BMC is about 10 – 550 times higher than the predicted C_ss_. For atrazine, the BMC is about 2 times higher than the predicted C_ss_. This direct comparison of *in vitro* derived BMC with *in vivo* C_ss_ could not identify the known hepatotoxicants as a potential hazard for humans.

As the C_ss_ metrics are not available for all chemicals, we tested the use of a three-compartment steady state PK model to derive oral equivalent doses (OED) from respective BMC values. OEDs for chemicals were predicted for chemicals with available exposure data such as human therapeutic doses, doses from clinical trials, food and drinking amounts, or experimental doses in animal models. OED was then predicted for every endpoint for which a significant response was observed in the imaging assay. The predicted OED values across all endpoints were then summarized into a single box plot per chemical (**Figure 7 and Table 4**). Predicted OED values overlapped with published human dose data for fialuridine, tetracycline, amiodarone, valproic acid and menadione, whereas OED values of Rotenone, WY14643, Lomitapide and Metformin were 10–10’000 times higher than realistic exposure levels. The OED values of orotic acid and carbosulfan overlapped with their reported no observable adverse effect level (NOAEL) and lowest observed adverse effect level (LOAEL) concentrations. Additionally, orotic acid’s OED prediction overlapped with the corresponding exposure estimate. The OED value for carbofuran was about 2,000 to 20,000 times higher than the published NOAEL/LOAEL, while for atrazine it was 10 – 100 times smaller than the published NOAEL/LOAEL values, and maximum residue levels (MRL) were 10–100 times lower. Finally, the predicted OED for carbosulfan was 2000 times higher than MRL.

**Table 4.** Comparison of predicted oral equivalent doses (OED) with published exposure values

| Chemical | OED<sup>1</sup> (mg/kg/d) | *In vivo* exposure<sup>2</sup> (mg/kg/d) | *In vivo* exposure reference |
| --- | --- | --- | --- |
| rotenone | 0.3 | 0.01 | (United States Environmental Protection Agency (USEPA), 2007) |
| fialuridine | 5.6 | 0.07 – 0.25 | (Kleiner *et al.*, 1997) |
| amiodarone | 15.8 | 2.9 – 22.9 | (Siddoway *et al.*, 2003) |
| tetracycline | 30.2 | 5.4 – 14.3 | (Bienenfeld *et al.*, 2017) |
| WY14643 | 1.7 | 0.4 – 0.7 | (Pollinger and Merk, 2017) |
| valproic acid | 92.7 | 3.6 - 30 | (Koch-Weser and Browne, 1980) |
| menadione | 55.4 | 1.1 – 36.6 | (Hu *et al.*, 1996) |
| lomitapide | 1333 | 0.07 – 0.86 | (Panno *et al.*, 2014) |
| metformin | 6778 | 7.1 – 35.7 | (Garber *et al.*, 1997) |

<sup>1</sup>In vitro to *in vivo* extrapolation derived median, <sup>2</sup>Exposure values include human therapeutic ranges or doses given to rat and rabbits

## 5 Discussion

In this study we developed a quantitative approach for simultaneous evaluation of chemicals for their potential to induce cellular alterations in nuclear morphology, lipid accumulation, mitochondrial membrane potential and oxidative stress as key hallmarks of steatosis etiology. Thus, a high content imaging assay was combined with PBPK modeling to derive relevant human dose predictions. Validation studies were carried out using 14 reference chemicals with an overall sensitivity and specificity of 63% (**Figure 8**). Notably, 6/9 predicted OED values for selected reference chemicals overlapped with estimated exposure values, supporting the relevance of this approach to risk assessment. Among FDA-approved and experimental drugs and food-related chemicals with chemical features associated with nuclear-receptor binding, one or more key events could be induced and quantified after 24 h exposure. Moreover, single cell analysis of tetracycline-exposed cells revealed that hepatocyte-like cells accumulated more lipid droplets and had increased MMP and oxidative stress levels compared to cholangiocyte-like cells, demonstrating the potential to extract cell type-specific effects in mixed cell populations. Finally, 14 food-related chemicals with structural alerts for receptor binding as a key initiating event in steatosis, were screened by this assay. Amongst the tested compounds and endpoints, orotic acid was identified to disrupt mitochondrial membrane potential, but not other markers and the predicted OED overlapped with estimated human exposure values. Novel aspects of the work reported here include the high accuracy of predicted OED with human exposure values for known chemicals and the application of the method to screen food-related chemicals.

**Figure 8.**
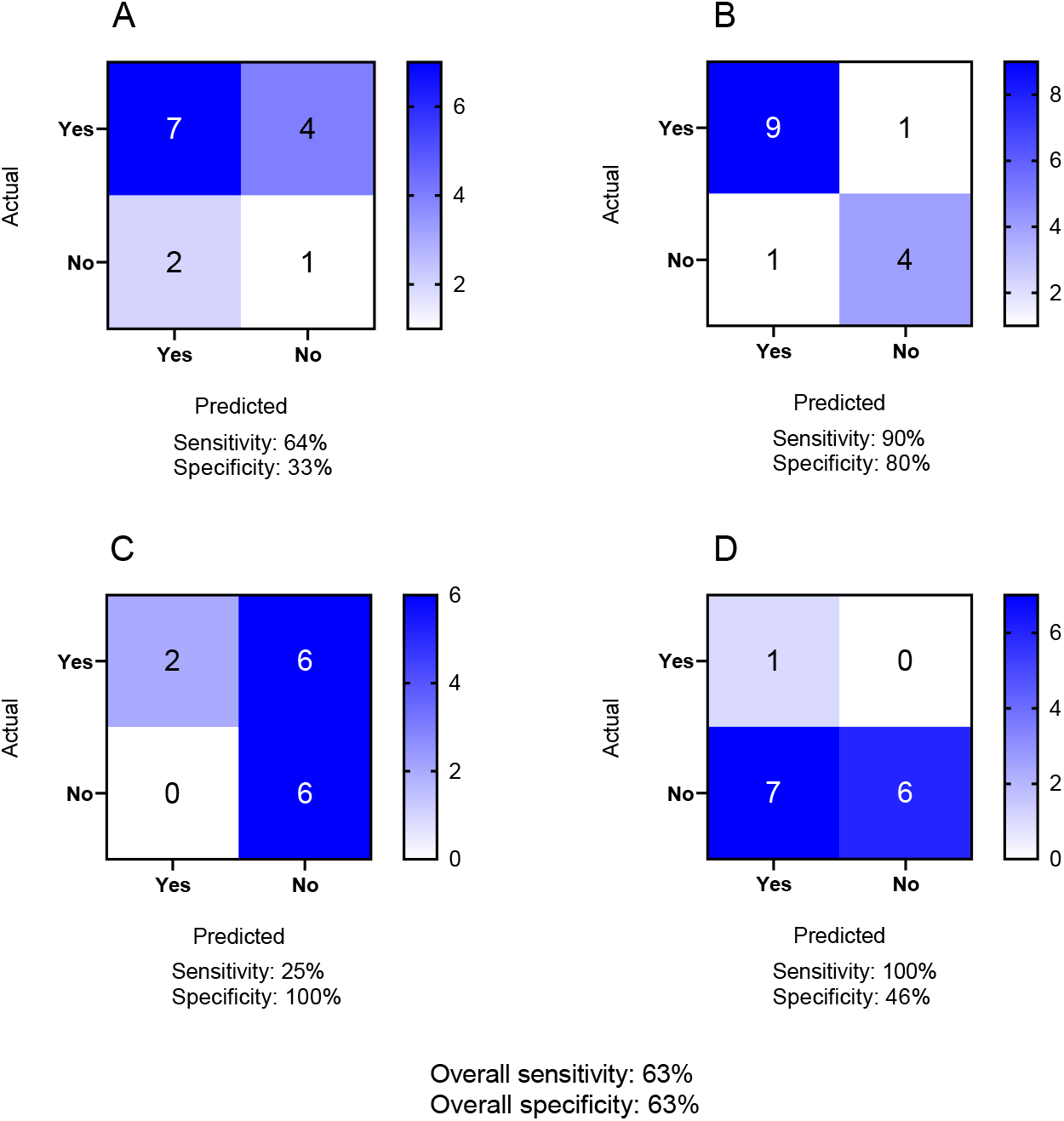
Quantification of the predictivity of the in vitro assay for A) lipid accumulation B) mitochondrial membrane potential C) oxidative stress and D) nuclear morphology for 14 reference chemicals using confusion matrices. Sensitivity was calculated as “number of true positives/(number of true positives + number of false negatives)” and specificity was calculated as “number of true negatives/(number of true negatives + number of false positives)”. Overall sensitivity and specificity were calculated across all 4 endpoints.

In this study we correctly flagged 7/11 chemicals known to induce lipid accumulation, and also identified two chemicals, metformin and WY-14693, as having the potential to induce lipid accumulation, that were unexpected (**Figure 8**). Potential rationale for this observation, in the case of metformin, is that it induced lipid accumulation only at extremely high, physiologically irrelevant concentrations (25’000 and 50’000 μM), whereas data suggesting metformin reduces lipid accumulation was observed for concentrations ranging from 100 μM to 2 mM (Kim *et al*., 2020; Zare *et al*., 2019). In addition, 4 chemicals, lomitapide, etomoxir, fialuridine and β-naphthoflavone did not give rise to lipid accumulation as we would have expected based on their established mechanism of action and previous clinical observations (**Table 2**) (McKenzie *et al*., 1995; Cuchel *et al*., 2013). A possible explanation for these apparent false negative results is that the 24-hour duration of exposure may be too short for these chemicals to induce lipid accumulation. Indeed, lipid accumulation in patients was observed only after 13 weeks for fialuridine and 26 weeks for lomitapide (McKenzie *et al*., 1995; Cuchel *et al*., 2013). This limitation suggests that future research using the assay reported here, but beyond the scope of the present work, should aim to evaluate how longer duration in vitro exposures, i.e. on the order of one to two weeks may resolve false negative results. Encouragingly, in previous studies hepatotoxicity was effectively characterized in HepaRG cells after up to 14 days of chemical exposure(Donato *et al*., 2022; Dietrich *et al*., 2020; Anthérieu *et al*., 2011).

In addition to lipid accumulation data, we could assess changes in mitochondrial membrane potential, oxidative stress and nuclear morphology with varying degrees of accuracy (**Figure 8**). Thus, in the case of the 10 chemicals expected to disrupt MMP, 9 were positive in our assay, whereas one chemical, valproic acid, did not affect MMP as we would have expected based on its mechanism of action (**Table 2**) (**Figure 8**). This is potentially expected since valproic acid has been characterized to induce mitochondrial dysfunction at very high concentrations, i.e. above 15 mM, and after prolonged exposure up to 72 h (Caiment *et al*., 2020) whereas we tested concentrations ranging from 94 to 6,000 μM and exposed the cells for 24 h. Interestingly, etomoxir and fialuridine increased MMP in a dose-dependent manner and exceeded the threshold at higher concentrations, which to our knowledge has not been reported previously. However, the biological relevance of this observation is still unclear and needs further investigation.

In the case of oxidative stress, sensitivity was poor as of the 8 chemicals expected to induce oxidative stress, only amiodarone and tetracycline actually did, six were unexpected negative including the well-known ROS-inducer menadione (**Figure 8**). This apparent low sensitivity could also be due to potential duration differences and a capacity for adaptive responses since menadione induced oxidative stress in embryo chick cardiomyocytes after exposure durations as short as 25 min (Loor *et al*., 2010). Thus, in addition to longer than 24 h durations mentioned previously, also shorter exposure durations than 24 h could be important to provide a more dynamic picture of oxidative stress responses in hepatocytes. Finally, nuclear morphology proved to be a highly or potentially over-sensitive read-out in this assay. Whereas only 1 chemical was expected to induce such changes (menadione), altered nuclear morphology was actually observed for a total of 8 chemicals. Nuclear morphology response mostly was above the calculated threshold at the highest tested concentrations, consistent with cell viability data (**Supplementary figure 4**).

After having validated the high content screening assay with reference chemicals, we applied the assay to data-scarce food-related chemicals. A majority of the food-related chemicals did not induce any cellular response however a salient outcome was the observation that orotic acid disrupted mitochondrial membrane potential. Furthermore, the extrapolated doses overlapped with actual exposure values (**Table 5**). Indeed, orotic acid is known to induce fatty liver in rats, whereas other animal species including mice and monkeys did not develop fatty liver upon orotic exposure (Durschlag and Robinson, 1980; Löffler *et al*., 2015). These findings are consistent with a concern that orotic acid might be a potential risk factor for steatosis in humans, however further investigations are needed.

**Table 5.** IVIVE of selected food-related chemicals which showed a response above the calculated threshold in the imaging assay.

| Chemical | LA<sup>1</sup> | MMP<sup>1</sup> | OS<sup>1</sup> | NM<sup>1</sup> | Median C<sub>ss</sub><sup>2</sup> (µM) | In vivo blood plasma conc. (µM) | OED<sup>3</sup> (mg/kg BW/day) | NOAEL/LOAEL from literature (mg/kg BW/day) | Exposure estimate from literature | Maximum residue levels (mg/kg) | Literature |
| --- | --- | --- | --- | --- | --- | --- | --- | --- | --- | --- | --- |
| orotic acid | - | 423 | - | - | 7.04 | 0.15 – 0.3 | 61.2 (95% CI 21.1 – 134) | 50 / 100 | 2 – 100 mg/kg BW/day | - | (D'Apolito *et al.*, 2012; Aguilar *et al.*, 2009) |
| uric acid | 591 | - | - | - | 6.84 | 217 – 402 | 87.3 (95% CI 67.3 – 150.6) | - | - | - | (Li *et al.*, 2009) |
| atrazine | 19 | - | - | - | 40 | - | 0.5 (95% CI 0.2 – 1.7) | 5/50 (increase relative liver weight) | - | 0.05 – 0.1 | (Agency for Toxic Substances and Disease Registry (ATSDR), 2003) |
| carbofuran | - | 4975 | - | - | 9 | - | 565 (95% CI 91.7 – infinity) | 0.03 – 0.1/0.1-0.3 | - | 0.01 – 0.5 | (Food and Authority, 2009a) |
| carbosulfan | 77 | - | - | - | 8.5 | - | 10.4 (95% CI 7 – 19.3) | 0.5 – 1.2/5 - 62 | - | 0.005 | (Food and Authority, 2009b) |

LA: lipid accumulation, MMP: mitochondrial membrane potential, OS: oxidative stress, NM: nuclear morphology, <sup>1</sup>Benchmark concentration (µM) per imaging endpoint calculated with BMDS software, <sup>2</sup>Derived from in vitro to in vivo extrapolation, <sup>3</sup>In vitro to in vivo extrapolation derived median

In vitro response data typically describes population average responses, missing potentially relevant information about how subpopulations might react to chemical exposure (Singh *et al*., 2014). In our study, therefore, we performed single cell analysis and investigated how hepatocytes and cholangiocyte-like cells responded on a single cell level to tetracycline exposure. For tetracycline in particular, we could readily observe from preliminary measurements that responses of individual cells were heterogeneous, so we quantified the distribution. To our knowledge, this is the first time that chemically exposed HepaRG cells were analyzed on a single cell level, discriminating cellular responses in hepatocyte and cholangiocyte-like cells. Interestingly, hepatocytes always responded more than cholangiocyte-like cells, possibly because hepatocytes may take up chemicals more efficiently than cholangiocyte-like cells do. Furthermore hepatocytes with larger lipid droplets were rather protected from a decrease in mitochondrial membrane potential compared to hepatocytes with small- and medium-sized lipid droplets. Likewise hepatocytes with larger lipid droplets tended to have lower oxidative stress levels than hepatocytes with smaller ones suggesting a protective effects of larger sized lipid droplets (Jarc and Petan, 2019). This kind of analysis can be used as a basis to formulate new hypothesis about mechanisms of toxicity.

PBPK models can be used to perform *in vitro* to *in vivo* extrapolation and predict possible human exposure scenarios that connect observed *in vitro* effects with human exposure levels via predicted OED values (Blaauboer, 2010). In our study, predicted Css values overlapped with Css values observed *in vivo* for all chemicals except BNF and WY14643 (**Figure 7A**). Importantly, for compounds with available human exposure data, estimated OEDs were within the range of published exposure values. Nevertheless, nominal in vitro concentrations for selected chemicals were higher compared to in vivo Css (e.g., metformin or valproic acid), leading to a high estimate of the OED. The following assumptions were made during IVIVE performed in this study: 1) Constant dose rate of 1 mg/kg/d with complete absorption for every chemical and excretion was limited to the renal route; 2) Plasma protein binding was not considered when in vitro assay concentrations were used for deriving BMC; 3) Metabolism of the chemicals was not accounted for. Aside from the quantitative aspects, there are also uncertainties associated with IVIVE regarding translation of whether cellular effects observed in vitro also occur in vivo, or whether adaptive or further detrimental responses at the organ level are relevant. Quantitative relationships between higher level key events in the AOP are needed to further link the cellular phenotypes with human disease. This would need to be further validated in more complex and physiologically relevant models and compared to relevant human exposures, including in susceptible populations, in order to inform risk assessment.

The approach presented here provides a new method combining high content imaging of chemically exposed human liver cells and IVIVE to prioritize chemicals for further evaluation as steatosis risk factors. Further validation of the specificity and sensitivity of the in vitro model and methodology by testing more extensive and diverse groups of chemicals (including mixtures of chemicals) is suggested for further study. The approach may also be adapted for more complex *in vitro* models such as liver spheroids from primary human cells (including Kupffer cells and hepatic stellate cells, in addition to hepatocytes) to potentially improve the prediction accuracy and biological relevance of the hepatic spheroid model to progressive NAFLD, which also involves persistent inflammation and fibrosis. These apical endpoint investigations could be combined with molecular mechanistic data derived from transcriptomic and proteomic analyses to identify additional predictive key events and refine mechanistic understanding. As a final step, the adaptation of this approach for high-throughput screening of existing chemicals of concern and of emerging chemicals would support a first step in a tiered approach addressing the needs of next generation chemical risk assessment for a disease of emerging human health relevance.

## 8 Acknowledgements

The authors acknowledge the financial support from the Swiss Federal Food and Veterinary Office and the support of the Scientific Center for Optical and Electron Microscopy ScopeM of the Swiss Federal Institute of Technology ETHZ. The authors would also like to thank Dr. Katherine Hurley and Shuhuan Zhai for their critical feedback during manuscript writing

