## Supplementary files for "Chemical screening of food-related chemicals for human fatty liver risk: Combining high content imaging of cellular responses with in vitro to in vivo extrapolation"

for

This supplementary information includes:

- Supplementary figures 1 – 5
- Supplementary tables 1 – 3

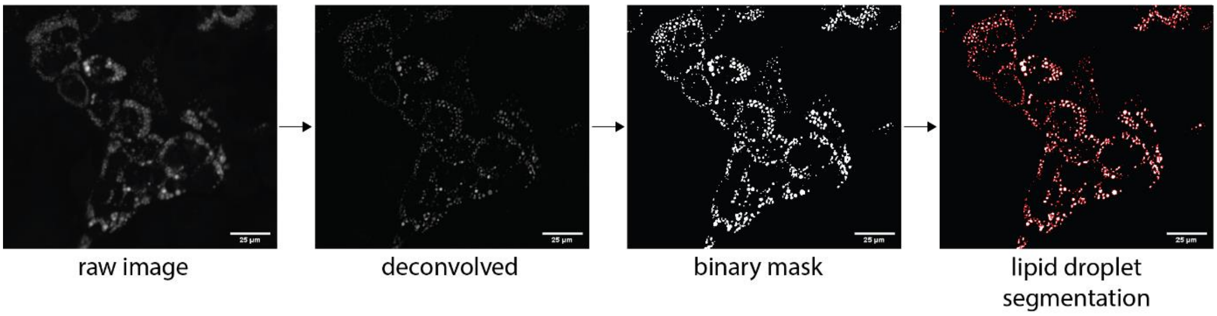

*Supplementary figure 1. Lipid droplet quantification. Input was a raw fluorescent image stained with Bodipy 505/515 to identify lipid droplets. Then background was substracted and deconvolved using Fiji. These images were then loaded into CellProfiler and a binary mask was created. Based on this binary mask, single lipid droplets can be segmented. Scale bar = 25 µm*

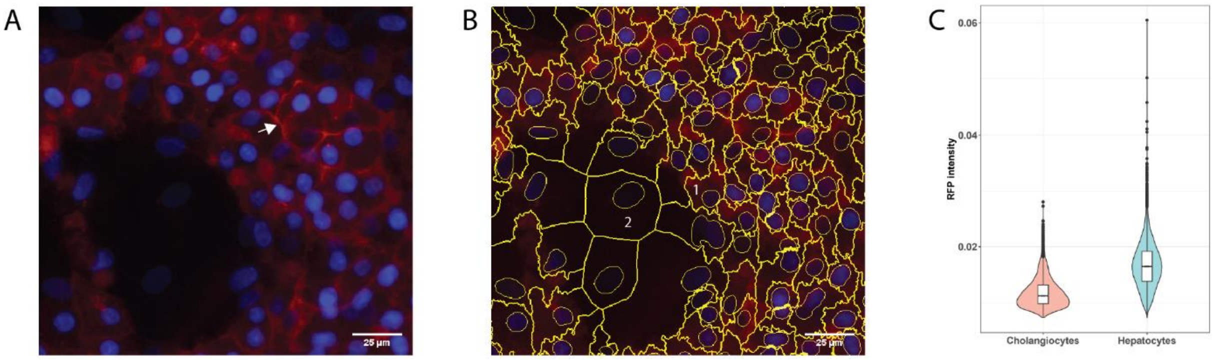

*Supplementary figure 2. Anti-ASGPR1 staining in HepaRG cells. A) Representative fluorescent image of HepaRG cells stained ASGPR1 (hepatocytes; red) and Hoechst (nuclei; blue). Staining is observed in hepatocyte island whereas cholangiocytes remain unstained. Membrane localization of ASGPR1 was visible (white arrow). B) An example image of segmented cells (yellow lines). Hepatocytes were identified using two criteria: (i) a smaller cytoplasm compared to cholangiocytes (ii) stained positive for ASGPR1. C) Single cell quantification of ASGPR1 intensity in hepatocytes and cholangiocytes using single cell analysis pipeline. Between 5000 and 10'000 cells were analyzed per cell type. Scale bar = 25 µm*

*Supplementary table 1: Physicochemical properties and in vivo toxicokinetic data for selected chemicals where Clint was manually calculated using formula 3 and added to the httk database*

| Name | CAS | MW (g/mol) | LogP | Fup | In vivo clearance (L/h) | Calculated Clint (ul/min/10^6 cells) | Species | Literature |
|---|---|---|---|---|---|---|---|---|
| Lomitapide | 182431-12-5 | 693.7 | 8.2 | 0.002 | 573.4656 (after 50 mg p.o.) | 48.25141 | Human | (Aegerion Pharmaceuticals, 2012) |
| Fialuridine | 69123-98-4 | 372.09 | -0.9 | 0.375 | Total clearance: 6.27 Renal clearance: 4.21 Nonrenal clearance: 2.06 | 0.1733285 | Human | (Bowsher *et al.*, 1994) |
| Metformin | 657-24-9 | 129.16 | -2.6 | 0.99 | Total clearance: 79.1 Renal clearance: 31.5 Nonrenal clearance: 31.5 | 4.005065 | Human | (Scheen, 1996) |
| Beta-Naphthoflavone | 6051-87-2 | 272.3 | 4.4 | 0.04 | 1.3 (plasma clearance) | 23.36091 | Rat | (Adedayo Adedoyin, Leon Arons, 1993) |
| Menadione | 58-27-5 | 172.18 | 2.2 | 0.2 | 49.32 | 58.33511 | Rabbit | (Hu *et al.*, 1996) |

Fup: fraction unbound plasma, Clint: in vitro intrinsic hepatic clearance

*Supplementary table 2: Z' and strictly standardized mean difference (SSMD) comparison of positive controls*

| **OAPA, LD** | | | | **Amiodarone, OS** | | |
|---|---|---|---|---|---|---|
| **Plate Nr.** | Z' | SSMD | | Plate Nr. | Z' | SSMD |
| **1** | 0.61 | 10.72 | | 1 | 0.19 | 4.25 |
| **2** | 0.7 | 13.87 | | 2 | 0.22 | 4.15 |
| **3** | 0.66 | 12.11 | | 3 | 0.26 | 4.63 |
| **mean** | 0.656667 | 12.23333 | | mean | 0.223333 | 4.343333 |
| **Rotenone, MD** | | | | **Menadione, NM** | | |
| **Plate Nr.** | Z' | SSMD | | Plate Nr. | Z' | SSMD |
| **2** | 0.147 | -4.7 | | 1 | 0.494 | 7.64 |
| **3** | 0.45 | -7.63 | | 2 | -0.065 | 3.19 |
| **4** | 0.12 | -4.33 | | 3 | 0.06 | 3.76 |
| **mean** | 0.239 | -5.55333 | | mean | 0.163 | 4.863333 |

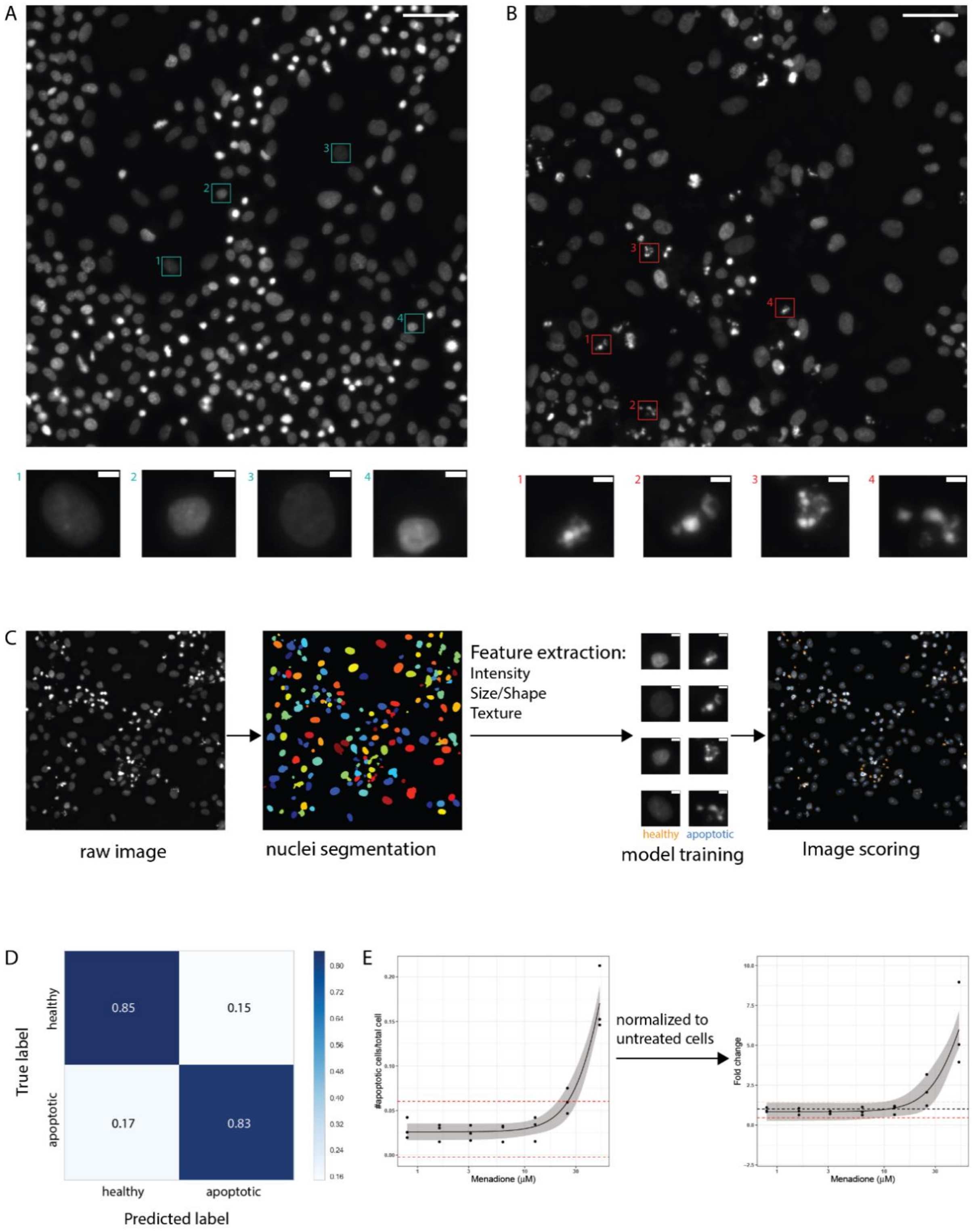

*Supplementary figure 3.Machine learning approach to quantify number of apoptotic cells. A) Representative fluorescent image of untreated HepaRG cells stained with Hoechst. In small boxes are examples of healthy nuclei. Scale bar in big picture is 25 µm and in the small boxes 5 µm. B) Representative fluorescent image of HepaRG cells treated with 50 µM Menadione and stained against Hoechst. In small boxes are examples of apoptotic nuclei. Scale bar in big picture is 25 µm and in the small boxes 5 µm. C) Description of machine learning pipeline. First, a raw nuclei image is segmented and over 200 features (intensity, size/shape and texture) per nuclei were extracted. This features were then fed into a random forest classifier model to train the model. Subsequently, images were scored with the trained model. D) Confusion matrix of a scoring run. E) Normalization approach of apoptotic nuclei quantification. Number of apoptotic nuclei were counted and normalized to cell number. Then they were normalized to untreated cells to get fold change.*

*Supplementary table 3: Food-related chemicals and pesticides selected for the application screen*

| Name | Tested concentrations (µM) | $EC_{50}$ (95% CI) in µM from cell viability assay | Use | Predicted activation of nuclear receptors |
|---|---|---|---|---|
| Alpha-terpineol | 125, 250, 500, 1000, 2000, 4000, 8000, 16000 (WST-1)<br>19.5, 39.1, 78.125, 156.25, 312.5, 625, 1250, 2500 (HCl) | 2565 (1853 – 3276) | Component of essential oils | none |
| Deoxycholic acid | 7.8125, 16.625, 31.25, 125, 250, 500, 1000 (WST-1)<br>2, 4, 8, 16.125, 33.25, 62.5, 125, 250 (HCl) | 246 (211 – 282) | Emulsifier and secondary bile acid | none |
| Orotic acid | 62.5, 125, 250, 500, 1000, 2000, 3000, 4000 (WST-1)<br>16.125, 31.25, 62.5, 125, 250, 500, 1000, 2000 (HCl) | NA | Mineral carrier in dietary supplements | THR |
| Uric acid | 7.8125, 16.625, 31.25, 125, 250, 500, 1000 (WST-1)<br>7.8125, 16.625, 31.25, 125, 250, 500, 1000 (HCl) | NA | Metabolite of purine nucleotides | None |
| Tartrazine | 16.625, 31.25, 125, 250, 500, 1000, 10000 (WST-1)<br>16.625, 31.25, 125, 250, 500, 1000, 10000 (HCl) | NA | Food dye | none |
| Fructose | 24, 98, 390, 1560, 6250, 25000, 100000, 400000 (WST-1)<br>39.1,39.1, 78.125, 156.25, 312.5, 625, 1250, 2500, 5000 (HCl) | 6.6 (1.1 – 156258) | Food component | none |
| Bisphenol A | 50, 100, 200, 225, 250, 400, 800, 1600 (WST-1)<br>1.6, 3.215, 6.25, 12.5, 25, 50, 100, 200 (HCl) | 237 (2.18 – 2.56) | Plasticizer and contact material | LXR, AHR, AR, ER, GR, PR, THR |
| Carbosulfan | 0.01, 0.1, 1, 10, 100, 500, 750, 1000 (WST-1)<br>0.01, 0.1, 1, 10, 100, 500, 750, 1000 (HCl) | NA | Insecticide | PPAR, AHR, AR, ER, GR, PR, THR, PXR |
| Vinclozolin | 0.00001, 0.0001, 0.001, 0.01, 0.1, 1, 10, 100 (WST-1)<br>0.00001, 0.0001, 0.001, 0.01, 0.1, 1, 10, 100 (HCl) | NA | Fungicide | FXR, AHR, AR, ER, GR, PR, THR |
| Mepanipyrim | 0.01, 0.1, 1, 10, 50, 100, 150, 300 (WST-1)<br>0.01, 0.1, 1, 10, 50, 100, 150, 300 (HCl) | NA | Fungicide | AHR, AR, ER, GR, PR, THR |
| Atrazine | 1.15, 2.3, 4.6875, 9.375, 18.75, 37.5, 75, 150 (WST-1)<br>1.15, 2.3, 4.6875, 9.375, 18.75, 37.5, 75, 150 (HCl) | NA | Herbicide | none |
| Carbofuran | 0.33, 1, 3, 9, 27, 90, 300, 1000 (WST-1)<br>7.8125, 16.625, 31.25, 125, 250, 500, 1000 (HCl) | NA | Insecticide | none |
| Fipronil | 0.512, 1.280, 3.2, 8, 20, 50, 125, 500 (WST-1)<br>0.225, 0.45, 0.9, 1.875, 3.75, 7.5, 15, 30 (HCl) | 21 (15 – 27) | Insecticide | PPAR, AHR, AR, ER, GR, PR, THR, PXR |
| Metazachlor | 0.33, 1, 3, 9, 27, 90, 300, 1000 (WST-1)<br>1.175, 2.35, 4.6875, 9.375, 18.75, 37.5, 75, 150 (HCl) | 183 (106 – 261) | Herbicide | none |

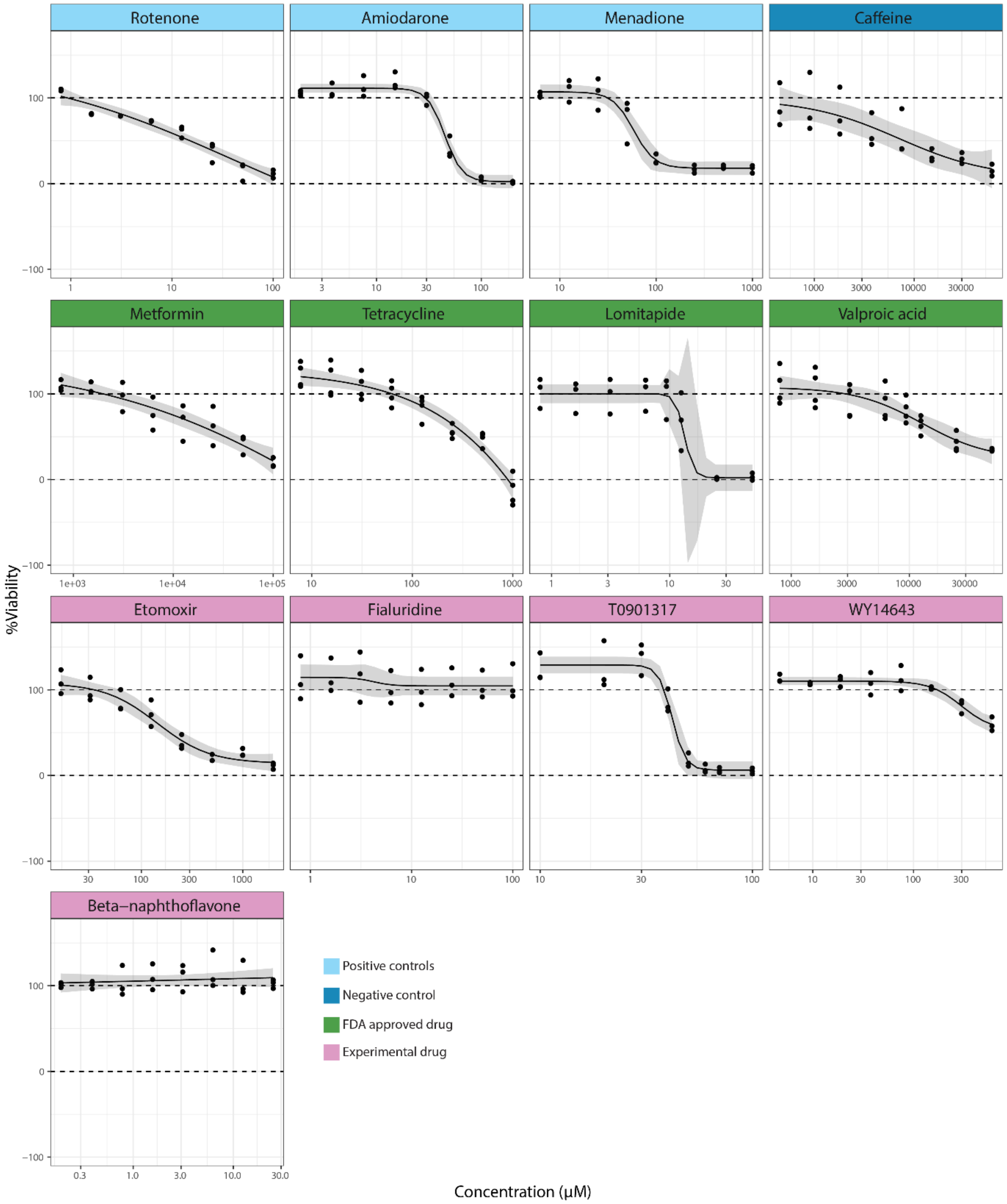

*Supplementary figure 4. Cell viability of reference chemicals assessed by WST-1 assay. Chemicals were grouped based on their usage as controls or current approval. Grey area around non-linear regression represents the 95% CI. Black dotted line represents a fold change of 1 (DMSO control). All data were first normalized to total cell number and then to untreated cells. At least 3 independent experiments with 6 technical replicates per concentration per experiment were conducted for all chemicals*

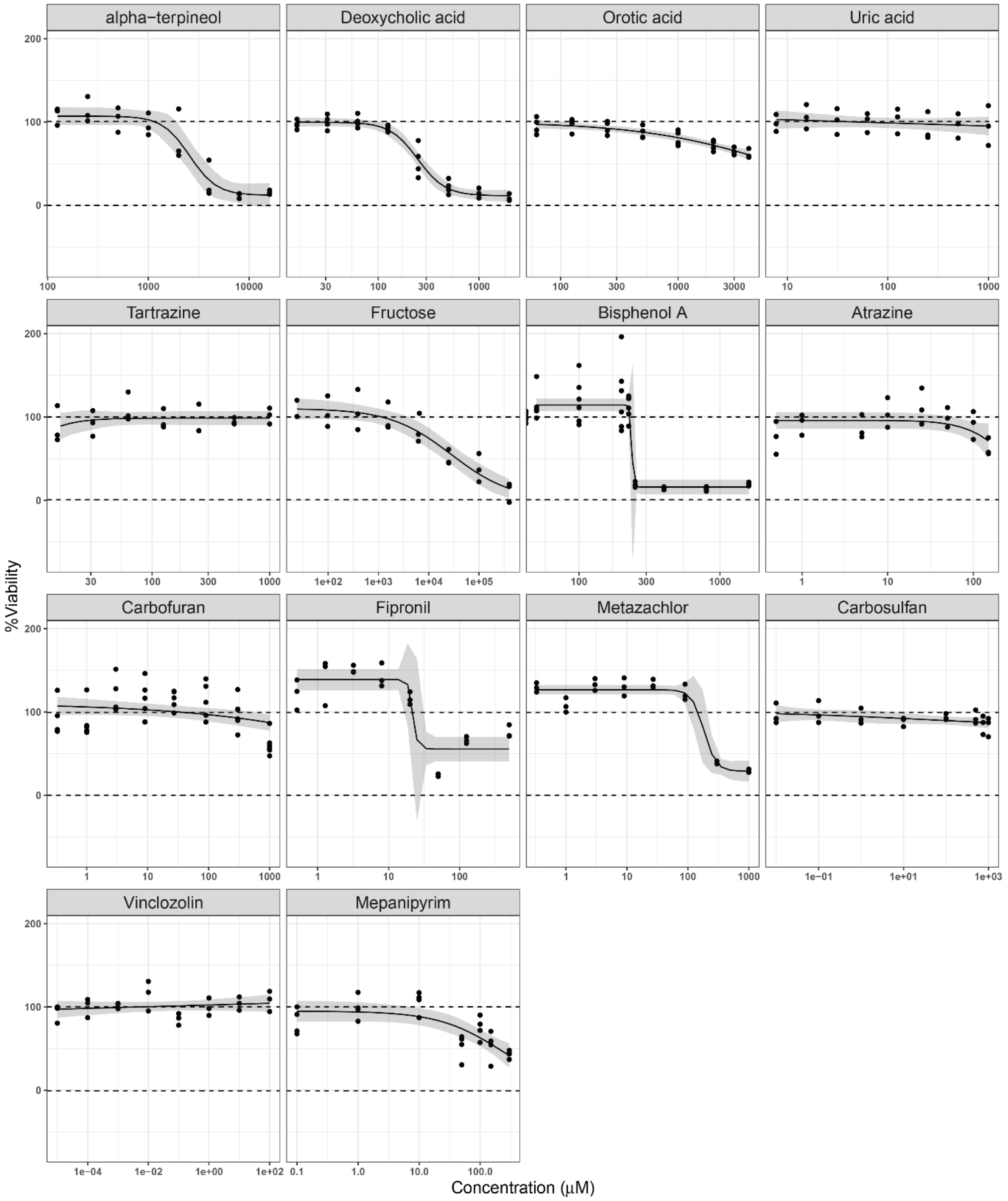

*Supplementary figure 5. Cell viability of food-related chemicals assessed by WST-1 assay. Grey area around non-linear regression represents the 95% CI. Black dotted line represents a fold change of 1 (DMSO control). All data were first normalized to total cell number and then to untreated cells. At least 3 independent experiments with 6 technical replicates per concentration per experiment were conducted for all chemicals*
